# Exploring the dynamics and interactions of the N-myc transactivation domain through solution NMR

**DOI:** 10.1101/2024.05.22.595265

**Authors:** Ewa Rejnowicz, Matthew Batchelor, Eoin Leen, Mohd Syed Ahangar, Mark W. Richards, Arnout P. Kalverda, Richard Bayliss

## Abstract

The myc family of proteins (c-, N- and L-myc) are transcription factors (TFs) responsible for maintaining the proliferative program in cells. They consist of a C-terminal domain that mediates heterodimerisation with Max and DNA binding, and an N-terminal disordered region culminating in the transactivation domain (TAD). The TAD participates in many protein–protein interactions, notably with kinases that promote stability (Aurora-A) or degradation (ERK1, GSK3) via the ubiquitin-proteasome system. Structural characterization of the TAD of N-myc, is very limited, with the exception of a crystal structure of Aurora-A bound to a helical region of N-myc. We probed the structure, dynamics and interactions of N-myc TAD using nuclear magnetic resonance (NMR) spectroscopy following its complete backbone assignment enabled by a truncation approach. Chemical shift analysis revealed that N-myc has two regions with clear helical propensity: one region within Trp77–Glu86 and the second between Ala122–Glu132. These regions also have more restricted ps–ns motions than the rest of the TAD, and, along with another known interaction site (myc box I), have comparatively high transverse (*R*_2_) ^15^N relaxation rates, indicative of slower timescale dynamics and/or chemical exchange. Collectively these features suggest differential propensities for structure and interaction, either internal or with binding partners, across the TAD. Solution studies on the interaction between N-myc and Aurora-A revealed a previously uncharacterised binding site. The specificity and kinetics of sequential phosphorylation of N-myc by ERK1 and GSK3 were characterised using NMR and showed no significant structural changes through the rest of the TAD. When doubly phosphorylated on residues Ser62 and Thr58, N-myc formed a robust interaction with the Fbxw7–Skp1 complex. Our study provides foundational insights into N-myc TAD dynamics and a backbone assignment that will underpin future work on the structure, dynamics, interactions and regulatory post-translational modifications of this key oncoprotein.

## INTRODUCTION

N-myc belongs to a family of transcription factor proteins which includes two other members, L-myc and c-myc. All three members are expressed in mammals and have functions in growth and development, but also have well documented pathological roles as oncoproteins in a wide variety of cancers [1–3]. The N-myc gene was discovered in a neuroblastoma cell line as a copy number amplified DNA sequence with sequence similarity to c-myc [4]. It is an important driver for cancers of neuronal origin such as neuroblastoma, medulloblastoma and retinoblastoma [5]. Consistent with this role as a driver oncogene, copy number amplification of N-myc is correlated with poor clinical outcomes in neuroblastoma [6–8]. The more extensively studied c-myc has been shown to act as a global amplifier of transcription, acting to increase transcription of already active genes [9–11]. c-myc has also been demonstrated to activate transcription of Pol I (ribosomal RNA), Pol II (mRNA), and Pol III (short, structured RNA) genes [9, 11–15]. However, at specific loci c-myc has also been shown to act as a transcriptional repressor, at least in relative terms [16, 17].

Myc proteins vary in size from 364 residues for L-myc to 464 residues for N-myc but have similar domain structures. The C-terminus of the protein contains a basic helix-loop-helix leucine zipper sequence of ∼90 residues, which forms a hetero-dimer with Max to produce a DNA binding domain that bind E-box sequences in the promoters and enhancers of genes [18–23]. The N-terminal region of the protein (∼370 residues in N-myc) is thought to be intrinsically disordered. This intrinsic disorder means that many of the functions of myc are brought about through recruitment of enzymes or structural proteins to sites close to E-box DNA sequences. Consistent with this role, mass-spec proteomics experiments have demonstrated that myc binds to hundreds of other proteins, either directly, or indirectly [24, 25]. There are six regions of conservation, known as myc boxes (MB), in the N-terminal region of the protein. Transgenic mice with N-myc replacing c-myc are both viable and reproduction competent [26]. This functional redundancy implies that the conserved sequences can mediate many, if not all, of the essential roles of myc. The transactivation domain (TAD) of myc transcription factors was defined in c-myc (residues 1–143) and contains the first three of these myc boxes (MB0, MBI, and MBII) [27]. Deletion of the first 150 residues of c-myc has been shown to abrogate both the capacity of c-myc to act as a transcriptional activator and to transform Rat-1a cells [28]. Recently, N-myc at elevated concentrations in neuroblastoma cells was shown to enter phase separated condensates in the nucleus, and that these condensates could differentially regulate gene expression. The TAD of N-myc (residues 1–137) was essential for this phase separation, further highlighting the importance of this part of the protein in myc-mediated oncogenesis [29].

Some of the specific functions of myc boxes and other sequences within the transactivation domain have been identified. MB0 (N-myc 18–37) and MBII (N-myc 110–126) recruit proteins such as histone acetyltransferase complexes (*e.g*., STAGA, TIP60), polymerase associated factors (*e.g*., General Transcription factor TFIIF), and other known transcriptional regulators (*e.g*., transcriptional intermediary factor 1α) [25]. In both c-myc and N-myc, the poorly conserved sequence between MBI (N-myc 45–63) and MBII is known to interact with molecules important for myc function. In c-myc, this region interacts with the General Transcription Factor complex, TFIID, part of the Polymerase II Pre-initiation complex [30]. In N-myc, this region interacts with Aurora-A kinase, which is an important complex for prevention of transcription–replication conflicts in S-phase [31, 32]. Recently a novel function of c-myc MB0 has been determined using NMR. MB0 binds to the DNA binding domain of the c-myc–Max dimer and appears to play a regulatory role in the complex binding to DNA [33]. MBI acts as a phosphodegron and is highly conserved across myc proteins; N- and c-myc have identical core MBI sequences (^47^EDIWKKFELLPTPPLSP^63^). Myc is phosphorylated at Ser62 either by ERK kinases, downstream of growth signaling, or by cell cycle kinases such as CDK1 or CDK2 [34–36]. Phosphorylation of Ser62 primes phosphorylation at Thr58 by GSK3 [34]. Thr58–Ser62 doubly phosphorylated species can bind to the Fbxw7 protein which is part of the of SCF^Fbxw7^ E3 ubiquitin ligase complex, this interaction results in ubiquitination of myc and subsequent proteasomal degradation [37–41]. In c-myc the MBI phosphodegron acts in concert with another phosphodegron around residue 244. An equivalent second phosphodegron has yet to be found for N-myc or L-myc [37]. This system is responsible for the short half-life of myc in cells, which for c-myc is 20–30 min [42].

Myc proteins are, for the most part, intrinsically disordered proteins (IDPs), they are regulated by post-translational modifications, interact with a large number of proteins, and tend to interact with these binding partners in the low micromolar range (*e.g*., Aurora-A 1 μM; GTFIIF 4.9 μM; TBP-TAF1 5.2 μM; Bin1 4.2 μM; PNUTS 3.5 μM) [25, 30, 31, 43, 44]. For these reasons nuclear magnetic resonance (NMR) spectroscopy is perhaps the most appropriate method to understand their structure, dynamics, regulation, and interactions. To a significant extent, previous NMR studies on myc have focused on the basic helix-loop-helix DNA binding domain and its interaction with Max [45–52]. The majority of backbone resonances for part of the c-myc TAD (1–88), which includes MB0 and MBI, have also been assigned [44]. This has been extremely useful to understand the dynamics, transient secondary structure, and effects of phosphorylation in this sequence. It has also been used to characterize the interaction between myc and binding partners such as PIN1 and Bin1 [44, 53]. Recently, backbone assignments have been determined for a large proportion of the c-myc backbone outside of the dimerisation and transactivation domains. The assignment was achieved using a divide and conquer approach using two polypeptides, c-myc 151–255 and 256–351 [33]. However, to date NMR has not been used to investigate the TAD of N-myc.

Here we present the first backbone NMR assignment of the N-myc transactivation domain. This includes the first myc assignment for MBII and the region known to interact with Aurora-A as a helix [31]. We characterize the dynamics of the molecule across timescales and orthogonally validate the presence of elements with latent secondary structure using circular dichroism spectroscopy. Additionally, we use NMR titrations to help explore the N-myc–Aurora-A complex, incorporating/implicating MBII as a further Aurora-A-binding region within an extensive interaction profile between the two proteins. Finally, we use the assignment to characterize the effects on N-myc structure of the MBI phosphorylation series and demonstrate that this material can form a complex with Fbxw7–Skp1.

## EXPERIMENTAL PROCEDURES

### Expression constructs

N-myc_1–137_ TAD and N-myc_64–137_ constructs were cloned into a pETM6T1 vector which contains a TEV-cleavable N-terminal His-NusA solubility/expression tag as described previously [54]. N-myc_18–59_ and N-myc_18–72_ were cloned into a pETM6T1 vector which contains the same TEV-cleavable N-terminal His-NusA tag followed by GB1 [55]. TEV cleavage leaves a short sequence GAM or GAMG in the constructs prior to Met1 in N-myc TAD, or Ser64 in N-myc_64–137_, respectively. N-myc_1–137_ TAD S7A was produced by site-directed mutagenesis using the Stratagene QuikChange protocol. TEV cleavage leaves a 62-residue sequence, including that of GB1, prior to Asp18 in GB1-N-myc proteins. The gene sequence of Fbxw7_261–706_ was cloned into a pET30-TEV vector with a TEV-cleavable 6x-His tag and the gene sequence of Skp1 was cloned into a pCDF-duet vector with no affinity tag. The Aurora-A kinase domain 122–403 C290A,C393A construct used was previously described [56].

### Protein production

Proteins were expressed using BL21 (DE3) RIL *E*. *coli* cells. Cells were transformed using the heat-shock method with the vector(s) of interest. 35 μg/mL chloramphenicol was used for routine maintenance of the RIL vector. 50 μg/mL kanamycin and 100 μg/mL spectinomycin was used to select and maintain pET and pCDF vectors respectively. A single colony was grown overnight in LB at 200 RPM and 37 °C. 10 mL of this starter culture was used to seed each litre of LB media. The cells were grown to mid-log phase. For routine expression, cells were then induced by addition of 0.6 mM ITPG and incubated overnight at 200 RPM and 20 °C. Cells were harvested by centrifugation at ∼6,000*g* for 20 min. Pellets were either stored at -80 °C or processed immediately. For NMR-labelled proteins, LB cultures were grown to mid-log phase in LB as described previously, but instead of being induced they were first pelleted at 2500*g* at 20 °C for 20 min. Cell pellets were resuspended in PBS buffer to remove residual LB. Bacterial cultures were then centrifuged again as described above. The resultant pellets were resuspended in 250 mL of minimal media (for each litre of LB), transferred to an autoclaved 2.5 L flask and incubated at 20 °C for a further 2 h at 200 rpm. Following this incubation period, overnight expression was induced with 0.6 mM IPTG. Cells were finally harvested as outlined previously. Minimal media consisted of 0.002 g/mL of ^15^NH_4_Cl and 0.01 g/mL of ^13^C glucose (or unlabelled glucose for ^15^N-only expression) in 50 mM Na_2_HPO_4_, 25 mM KH_2_PO_4_, 20 mM NaCl supplemented with 2 μM MgSO_4_, 0.2 μM CaCl_2_, 0.01 mM FeSO_4_.7H_2_O, micronutrients and a vitamin solution (BME vitamins 100x solution, Sigma-Aldrich). The solution was syringe filtered (0.2 μm) prior to use.

### Protein purification

To purify N-myc_1–137_ and N-myc_64–137_, bacterial pellets were resuspended using TBS-TCEP buffer (25 mM Tris, 2.7 mM KCl, 137 mM NaCl, 2 mM tris(2-carboxyethyl)phosphine (TCEP), pH 6.9) supplemented with c0mplete^TM^ Mini EDTA-free Protease Inhibitor Cocktail tablets (Roche) and 10 mg lysozyme. Cells were lysed by sonication and then clarified by centrifugation at 50,000*g* for 1 h. Protein was initially purified using affinity chromatography using HIS-Select Cobalt Affinity Gel (Sigma) on a gravity flow column. TEV protease was added, and the solution was dialysed in TBS-TCEP buffer at 4 °C overnight. Dialysed protein was further purified by ion-exchange chromatography using a HiTrap-FF Q column (GE Healthcare), resulting in the elution of N-myc proteins in the flow-through and at low salt concentrations. Fractions containing N-myc were subjected to a Cobalt affinity subtraction step to remove His-tagged TEV protease. Finally, N-myc proteins were polished using size-exclusion chromatography (SEC) using a Superdex 16/600 S75 column (GE Healthcare). GB1-tagged N-myc proteins were purified in a similar manner with a few alterations: 25 mM MES, 200 mM NaCl, 5 mM β-mercaptoethanol, pH 6.5 was used instead of the TBS–TCEP buffer. The dialysis buffer was the same with the exception that the NaCl concentration was 150 mM. Ion exchange was not performed on these proteins. The final size exclusion buffer contained 20 mM (K/H)_3_PO_4_, 150 mM NaCl, pH 6.5.

Fbxw7 and Skp1 were purified using Ni-affinity chromatography followed by dialysis with TEV protease to cleave the His-tag on Fbxw7. The cleaved tag was removed by applying the dialysed protein again to the Ni-column. Fractions containing purified complex were concentrated and subjected to size exclusion chromatography using a Superdex 16/600 S200 column equilibrated in 20 mM Tris, 200 mM NaCl, 2 mM β-mercaptoethanol, 10% glycerol, pH 8.

The kinase domain of Aurora-A (Aurora-A 122–403, C290A, C393A) was purified as described previously [56]. Protein concentrations were measured by absorbance at 280 nm using extinction coefficients determined using the Expasy ProtParam tool (https://web.expasy.org/cgi-bin/protparam/protparam). Purified proteins were concentrated, aliquoted and used immediately or snap frozen in liquid nitrogen and stored at -80 °C prior to use.

### NMR spectroscopy

The majority of NMR spectra were recorded on a 750-MHz Oxford Instruments magnet equipped with a Bruker Avance console and TCI cryoprobe. Additional experiments were recorded on a 950-MHz Bruker Ascend Aeon spectrometer equipped with a TXO cryoprobe and a 600-MHz Oxford Instruments magnet equipped with a Bruker Avance console and a QCI-P-cryoprobe (a list of experiments is given in supplementary information Table S1). ^1^H–^15^N HSQC spectra were recorded between 10 and 37 °C, and peak positions tracked to transfer assignments across temperatures. Although variable across the sequence, intensities for the majority of peaks dropped at higher temperatures (Fig. S1B), so, coupled with a desire to prolong the lifetime of the sample, assignment spectra were recorded at 10 °C. The data were processed into spectra using NMRPipe/NMRDraw [57]; peak assignments and further analysis were carried out with CCPNmr Analysis (version 2.5) [58]. For calculations of secondary shifts (Δδ) , reference coil values generated using a web server hosted by the University of Copenhagen (https://www1.bio.ku.dk/english/research/bms/sbinlab/randomchemicalshifts2/ [59]) were subtracted from measured shift values. Further structural analysis using all measured shifts was carried out using the TALOS-N web server (https://spin.niddk.nih.gov/bax-apps/nmrserver/talosn/ [60]). Chemical shift perturbations (CSPs) were calculated using a 0.15 weighting for ^15^N shifts compared to ^1^H shifts.

^15^N longitudinal (*R*_1_) and transverse (*R*_2_) relaxation rates and ^15^N–^1^H heteronuclear NOEs were measured for resolved ^1^H–^15^N peaks of a 150 μM sample of N-myc TAD using the 600 MHz instrument. Spectra were recorded at 10 °C. The recycle delays were 2.5 s for *R*_1_ and *R*_2_ experiments and 5 s for the heteronuclear NOE experiment. Eleven relaxation periods ranging from 10 to 1600 ms were used for the *R*_1_ experiment, and twelve relaxation periods from 17 to 201 ms were used for the *R*_2_ experiment. In both cases, two of the relaxation periods were duplicated to facilitate error estimations. Peak intensities measurements and analysis of relaxation rates was performed using PINT [61, 62]. Some residues were excluded from these analyses due to peak overlap or where it was not possible to confidently follow peak intensity changes across the experiment.

Phosphorylation of ^15^N-labelled N-myc TAD was tracked by collection of ^1^H–^15^N HSQC spectra prior to and subsequent to addition of kinases [63]. Data were acquired on the 600 MHz spectrometer, equipped with a QCI-P-cryoprobe. TBS–TCEP buffer was supplemented with 4 mM MgCl_2_ and 1 mM ATP, and a concentration of 120 μM N-myc TAD was used. ERK1 and GSK kinases were purchased from the MRC Protein Phosphorylation Unit, University of Dundee, UK. Small aliquots (∼10 μL) were added step-wise to give a final kinase concentration of ∼300 nM. By way of control, there was no sign of phosphorylation of N-myc TAD by Aurora-A or by PLK1 kinases.

For NMR interaction studies, N-myc constructs and Aurora-A kinase domain (122–403, C290A, C393A) were separately buffer exchanged into the following buffer (20 mM (K/H)_3_PO_4_ pH 6.5, 150 mM NaCl, 2 mM β-mercaptoethanol, 1% glycerol, 5 mM MgCl_2_, 5 mM ADP). Buffer exchange was achieved by sequential dilution and concentration in centrifugal concentrators. Initial N-myc protein concentrations were 150–315 μM. ^1^H–^15^N HSQC peak positions were very similar to those in the original buffer so assignments were straightforwardly copied across. Aliquots of Aurora-A (305–400 μM were added up to a [N-myc]:[Aurora-A] molar ratio of 1:1.4.

### Circular dichroism spectroscopy of N-myc peptides

The following 17-residue N- and C-capped peptides were purchased from Biomatik Corp., Canada:

Ac-PGEDIWKKFELLPTPPL-NH_2_ (N-myc 45–61, MBI); Ac-EPPSWVTEMLLENELWG-NH_2_ (N-myc 73–89, AIH); Ac-GFSAREKLERAVSEKLQ-NH_2_ (N-myc 119–135, MBII+); Ac-GFSAAAKLVSEKLASYQ-NH_2_ (c-myc 137–153, MBII+). Peptides were initially dissolved to give 0.5–1.0 mM stock solutions in buffer composed of 1 mM sodium borate, 1 mM sodium citrate, 1 mM (Na/H)_3_PO_4_ and 10 mM NaCl, pH 7 [64, 65]. For measurements, samples were further diluted in buffer to concentrations of 10–100 μM. Equivalent concentration solutions were also prepared containing 15 or 30% (v/v) trifluoroethanol. Spectra were recorded using ∼200 μL samples in 1 mm pathlength quartz cuvettes and an APP Chirascan CD spectropolarimeter. To maximise the secondary structural content, spectra were recorded at 5 °C. Data were collected every 1 nm for wavelengths 260–180 nm (1 nm s^−1^); each spectrum was recorded in duplicate. Due to significant Cl^-^ absorption, measurements <190 nm were unreliable and are not included. Background spectra were recorded using buffer alone, buffer/15% TFE or buffer/30% TFE, and background ellipticity values were subtracted from raw sample ellipticity values (*θ* (*λ*)) to calculate MREs using:

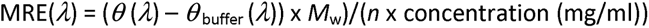

where *M*_w_ is the peptide molecular weight, and *n* is the number of total peptide bonds, in each case taken to be 17 accounting for the acetyl capping group. Helicity values were estimated using the following equation:

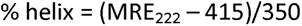

with MRE_222_ in units of deg cm^2^ dmol^−1^ res^−1^ [64, 66].

### Analytical size-exclusion chromatography

Analytical SEC was used to investigate the binding of N-myc TAD to the Fbxw7–Skp1 complex. Sample phosphorylation was achieved by addition of 1 mM ATP, 4 mM MgCl_2_, and 0.3 μM ERK1 and/or GSK3, and followed by NMR, and then samples were flash frozen to prevent further reaction. Prior to analytical SEC, N-myc TAD, in different phosphorylation states, was mixed in equivalent molar ratio with recombinant Fbxw7–Skp1 to a total volume of 100 μL. The proteins were incubated by slow rotation at 4 °C for 2 h prior to SEC. SEC was performed using a ÄKTA pure FPLC system and a Superose 12 10/300 GL column. The loop and sample volumes were 60 μL and 100 μL respectively. The flow rate was 0.5 mL/min. Fractions were collected every 0.5 mL and analysed using SDS-PAGE.

## RESULTS

### Backbone assignment of N-myc TAD

The primary amino acid sequence of N-myc_1-137_ (or N-myc TAD) reveals an amino acid composition typical for an IDP (Fig. 1) [67]. For example, the sequence has a comparatively large proportion of Pro (12%), Gly (10%) and Ser (10%) residues, which all disfavour secondary structure formation [68]. The sequence is also enriched in Glu (10%) and Asp (7%) residues, as is typical for transactivation domains [69], but has a low number of Ala (4%), Ile (2%) and Val residues (2%).

**Figure 1.**
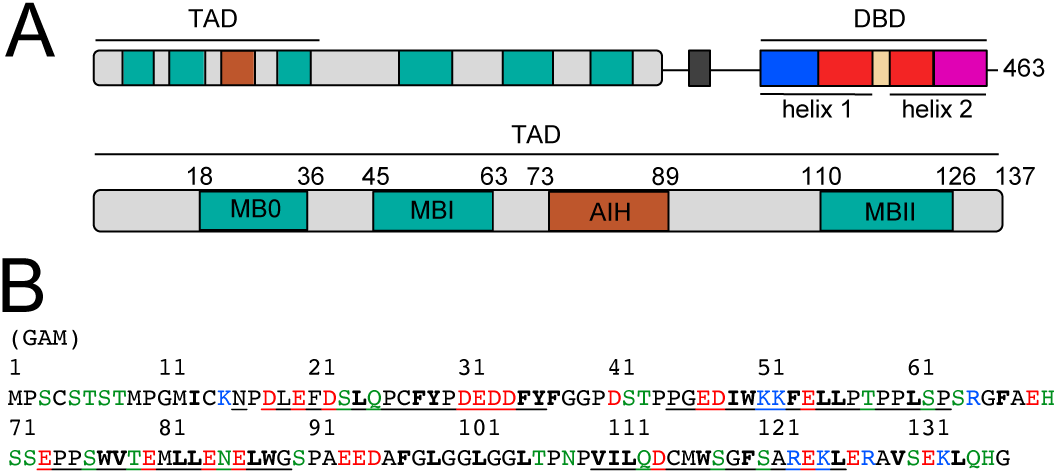
(A) Domain structure of N-myc showing the positions of the structured C-terminal DNA-binding domain (DBD), myc boxes and the N-terminal transactivation domain (TAD). The TAD is magnified below, showing the positions of myc boxes MB0, MBI and MBII and the Aurora-A interaction helix (AIH). (B) Primary sequence of the N-myc TAD; hydrophobic residues are shown in bold, basic residues in blue, acidic residues in red, polar residues in green. The positions of the myc boxes are underlined.

The ^1^H–^15^N HSQC spectrum of N-myc TAD (Fig. 2) shows the low dispersion characteristic of an IDP [70], but uncharacteristically—and as previously observed for c-myc—peaks are not uniformly sharp, exhibiting a range of line-widths, with a subset of broad peaks requiring an increased number of scans to be clearly observed. Usually, IDPs have fast rotational correlation times, resulting in slow *R*_2_, which is realised in the form of sharp peaks. The broad peaks indicate that there are regions that exhibit structure or otherwise altered dynamics within the N-myc TAD sequence (Fig. S1, particularly Lys51–Leu56, Ser76–Gly89 and Ile111–Ala122). This feature was initially probed by collecting spectra over a range of temperatures (283.1–310.2 K). Most peaks diminish in intensity as the temperature was raised; presumably increased solvent exchange outweighs the benefits of faster tumbling at higher temperatures. Some small regions (close to residue 30, 80 and 110) show increased intensities at moderate temperatures and are slower to lose relative intensity at high temperatures (Fig. S1B). Plotting chemical shift perturbations (CSPs) as a function of temperature (Fig. S1C) identified two regions, one close to Thr58 and the other near Ile111, that stand out as having large temperature coefficients. Both regions have a number of neighboring prolines. Residues in the region 120–130 have particularly low temperature coefficients, consistent with a level of protection afforded to the H–N bond through hydrogen bonding. A number of peaks for residues in this region moved in the opposite direction (downfield) on increasing temperature compared to peaks for the rest of the protein sequence (which uniformly moved upfield in ^1^H and ^15^N).

**Figure 2.**
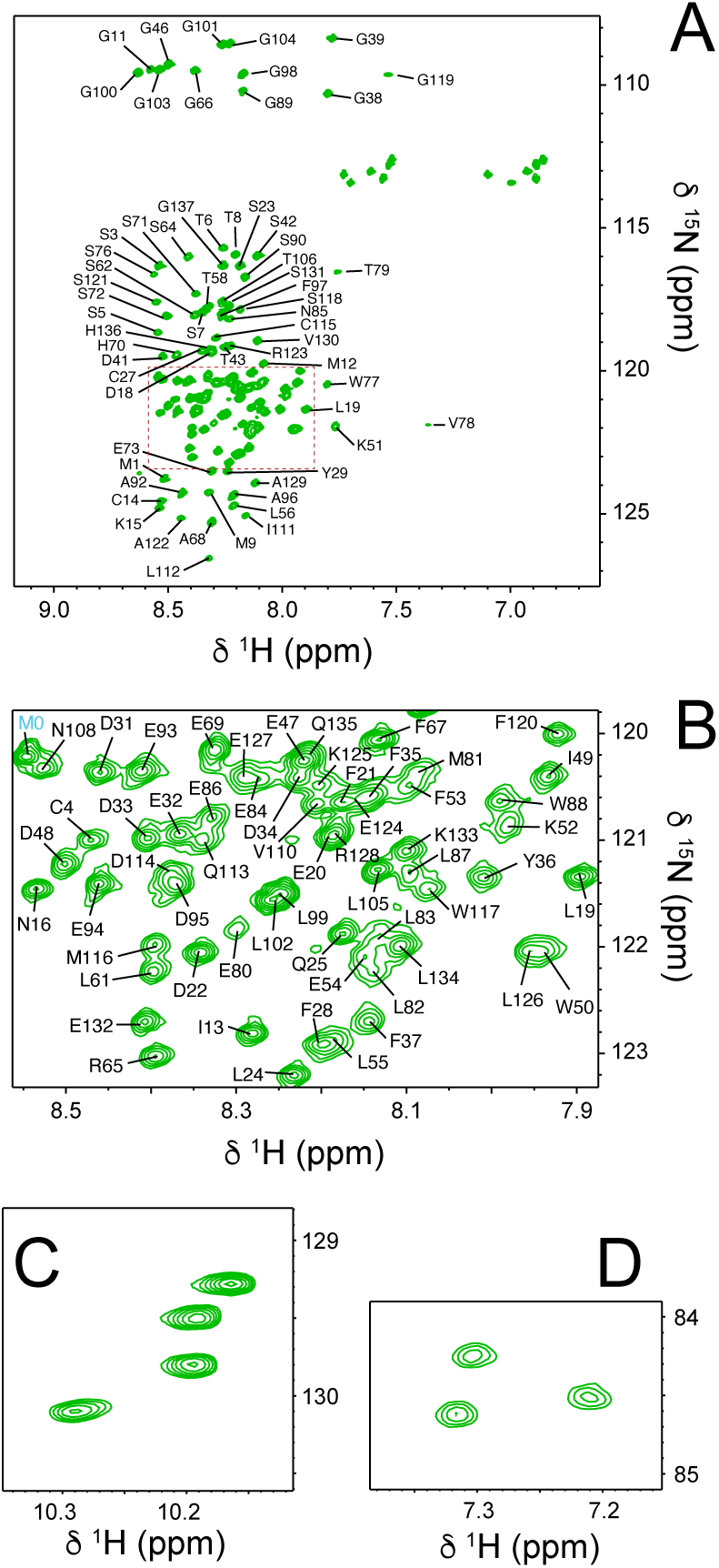
Assigned ^1^H–^15^N HSQC spectrum of the N-myc TAD. (A) Overall spectrum (omitting Trp and Arg Hε–Nε correlations). Plot shows the assignment of peaks outside the central region, note some of the weak outlying peaks *e.g*. Val78, Leu112, Lys51. Gln (ε) and Asn (δ) sidechain NH_2_ peaks (^15^N ∼ 113 ppm) are unassigned. (B) A magnified view of the central region showing the lack of dispersion here as well as variation in the intensity of peaks. (C, D) Sidechain Hε–Nε correlations. Four Trp (C) and three Arg (D) sidechain correlations are resolved (but unassigned) accounting for Trp51, Trp77, Trp88, Trp117, Arg65, Arg123, and Arg128.

Usefully, as a marker of protein integrity, the appropriate number of resolved sidechain resonances were observed for the four Trp Nε–Hε, six Gln (Nε–Hε) or Asn (Nδ–Hδ) pairs and even the three Arg Nε–Hε peaks (Fig. 2). N-myc TAD is labile to proteolysis, despite efforts made to purify the sample and use of protease inhibitors, as shown by the emergence of degradation peaks in the ^1^H–^15^N HSQC spectrum within around 3 days with the sample held at 10 °C.

The utilisation of standard BEST versions of ^1^H-detected 3D experiments (HNCO, HNcaCO, HNcoCA, HNCA, HNcocaCB, HNcaCB, HBHAcoNH, Table S1) enabled the unambiguous assignment of ∼60% of the N-myc TAD backbone. A few regions however were particularly challenging to assign; for example, residues Ser76–Trp88 were characterised by peak broadening and most peaks were positioned within regions of high ^1^H degeneracy within the ^1^H–^15^N HSQC spectrum causing peak overlap. Full assignment was achieved by recording assignment spectra for truncated sections of N-myc TAD and transferring assignments to the full-length construct to remove ambiguities. A C-terminal truncation of the TAD, N-myc_64–137_, and a fusion of GB1 [55] to N-myc_18–59_, provided excellent overlap with full length ^1^H–^15^N HSQC spectra for peaks away from the termini (Fig. S2). The shorter constructs yield spectra with fewer N-myc peaks and exhibit more favourable dynamics [71], and thus spectra have enhanced sensitivity and resolution in comparison to N-myc TAD spectra. There are the additional peaks from GB1 in the truncated fusion, but these have greater dispersion and generally do not interfere with the N-myc peaks. Peak position differences are also observed close to the termini in both cases, as would be expected. Small uniform differences in peak resonant frequencies, especially for pH-sensitive His residues, could be due to small variations in buffer. A suite of three 2D ^13^C-detected spectra (CON, haCACO and haCAnCO) and one ^15^N-detected spectra (haCAN) which avoid reliance of the congested ^1^H dimension were also collected for the N-myc TAD construct (Table S1) [72]. This corroborated the assignment and provided additional ^15^N, ^13^CO and ^13^Cα shifts for proline residues. All told, assignments for all backbone N, CO and Cα resonances within N-myc TAD were achieved; see BMRB entry 52066. Only 3 Cβ, 21 Hα and 23 Hβ shifts are not recorded.

### N-myc TAD structure and dynamics

Secondary chemical shift (Δδ) data can be used to indicate structural propensities within proteins [73]. For example, ^13^Cα Δδ values are positive in α-helices and negative in β-strands, whereas ^13^Cβ Δδ values display the opposite trend. In N-myc TAD (Fig. 3A), the close-to-zero ^13^Cα Δδ values and weak correlations between adjacent residues indicate that the N-terminal portion of N-myc TAD (aa 1–76) is largely random coil albeit with potentially some weak extended/β-strand propensity in MB0 (Leu24–Asp34) and weak helical propensity in MBI (Asp48–Phe53). There are two regions in the C-terminal portion of N-myc TAD (Trp77–Glu86 and Ala122–Glu132) with consistent positive values of ^13^Cα Δδ indicating that these parts of the protein have helical propensity. This pattern of ^13^Cα secondary shifts is very similar for the truncated GB1-N-myc_18–59_ and N-myc_64–137_ constructs, with the weak strand propensity in MB0 reproduced and consecutive positive secondary shift values shown between Val78–Asn85 and Ala122–Glu132 (Fig. S3A). The program TALOS-N [60] was used to provide a more general prediction of secondary structure utilizing more fully the backbone chemical shifts along the sequence (Fig. S3B). As expected, most of N-myc TAD is random coil but again TALOS-N places two α-helices at Trp77–Glu84 and Ala122–Glu132.

**Figure 3.**
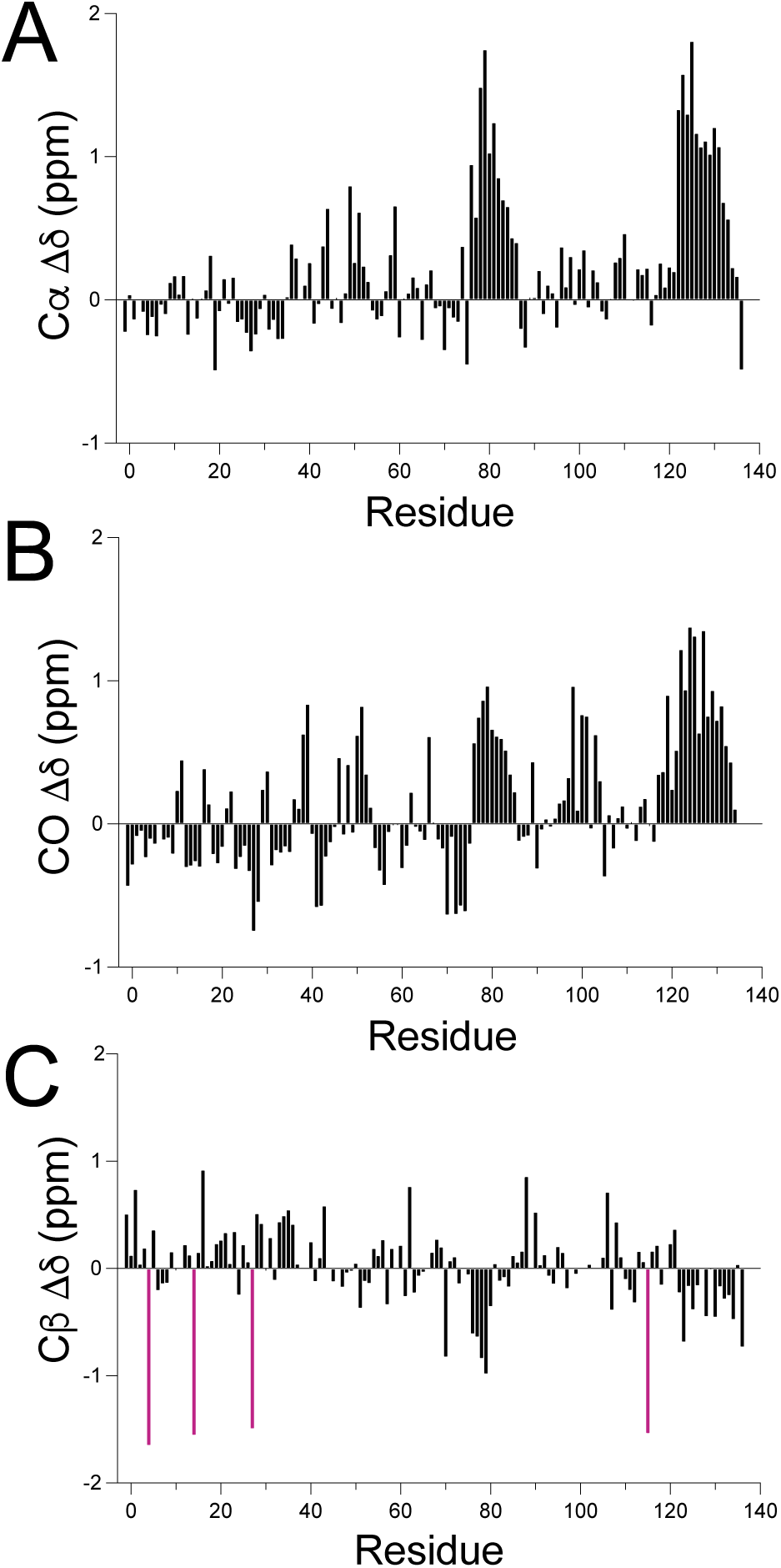
Structural propensities within N-Myc TAD. Secondary shifts (Δδ) for Cα (A), backbone carbonyl (B) and Cβ (C) nuclei across the sequence of N-myc_1-137_. Positive runs of Cα/CO Δδ values indicate α-helices, negative runs of Cα/CO Δδ values indicate β-strands and values nearing zero or with variation in sign of consecutive values indicate a lack of structure. The trends are mirrored for Cβ Δδ values. A comparison of Cα Δδ for N-myc TAD with the truncated variants is shown in Fig. S3A. The four Cβ Δδ values in purple are for the four Cys residues in the sequence; we contend that the anomalous values are due to an error in the reference coil value for Cys (Hendus-Altenburger, *et al*. (2019)).

To investigate further the behaviour of the N-myc TAD in the context of local and global structure, a set of three relaxation measurements, ^1^H–^15^N NOE, *R*_1_ and *R*_2_, was employed to probe the local dynamics of the protein. ^1^H–^15^N NOE values inform whether individual residue dynamics on the picosecond to nanosecond (ps–ns) time scale are determined by the overall rotational of the molecule (τ_C_) and are thus ‘fixed’ or rigid in nature, or whether their dynamics is faster than τ_C_, thus indicating a degree of flexibility [74]. Negative NOE values indicate large amplitude motions on short timescales as observed at the N- and C-termini of N-myc TAD (Fig. 4A) which have the highest degree of freedom in their motions [75]. Regions with restricted local movements result in increasingly positive NOE values as their rotational motions approach τ_C_. Most residues in N-myc TAD are characterised by ^1^H–^15^N NOE values in the range 0.2–0.3, similar to those observed in c-myc [44], indicating that most of the TAD is disordered. Within the TAD sequence there are stretches of residues which are characterised by higher NOE values (>0.35). These regions (Trp50–Phe53, Trp88–Ser90 and Ser118–Ser121) are likely to experience more restrictive motions of the N–H vectors (Fig. 4A) [76].

**Figure 4.**
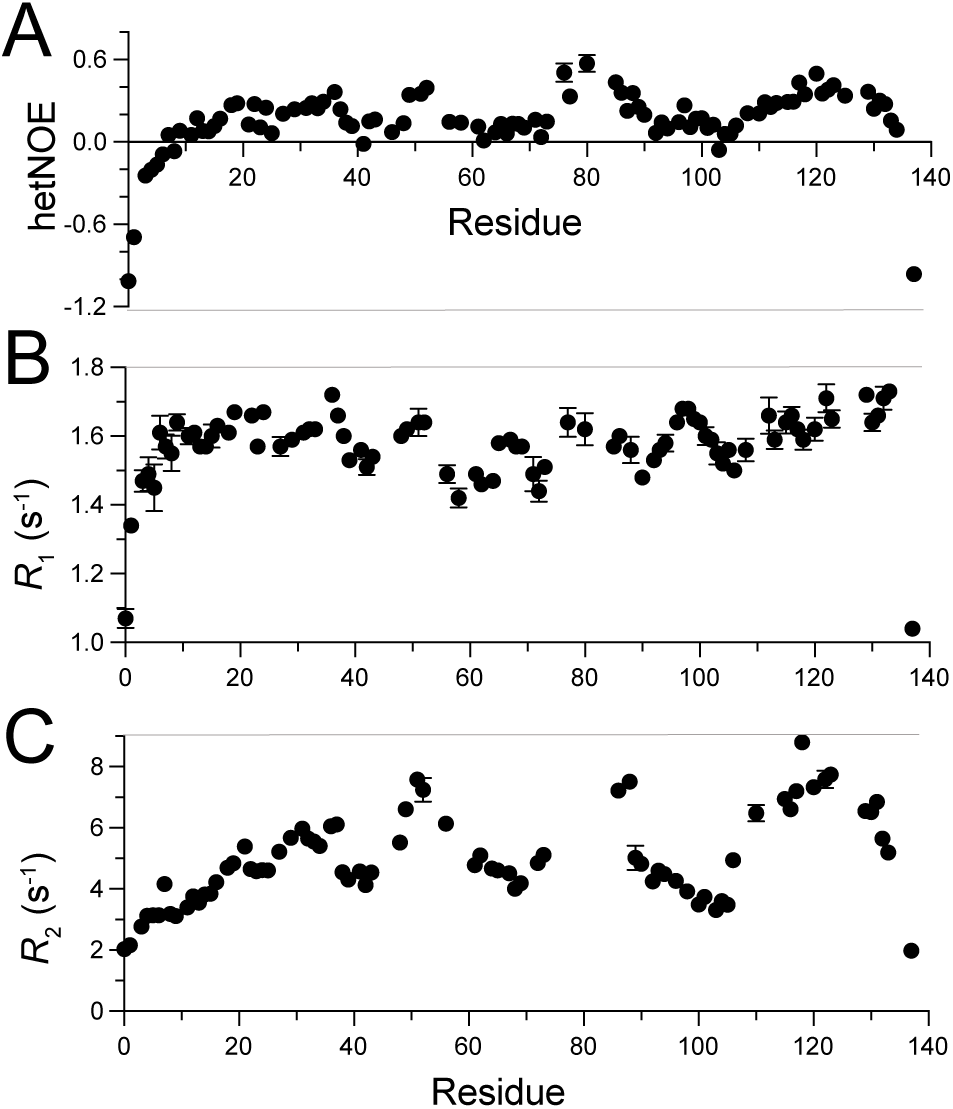
Relaxation measurements for the N-myc TAD. ^1^H–^15^N heteronuclear NOEs (A), longitudinal (*R*_1_, B) and transverse relaxation rates (*R*_2_, C). Experiments were carried out at 10 °C. Residues missing from the plots include prolines and overlapping/broad peaks for which intensity changes could not be accurately gauged.

*R*_1_ and *R*_2_ measurements (longitudinal and transverse relaxation rates, respectively) also report on ps–ns motions while *R*_2_ is also impacted upon by μs–ms time-scale changes [74]. Away from the termini, *R*_1_ values remain relatively consistent (1.4–1.7 s^-1^, average 1.6 s^-1^) across the length of N-myc TAD, with some subtle variation mirroring the hetNOE profile (Fig. 4B). This indicates that on the ps–ns time scale residues are broadly similar in their dynamics. *R*_2_ values increase with slower tumbling and are impacted by μs–ms dynamic processes, which signifies more ‘global’ conformational motions such as protein folding, oligomerisation, or chemical exchange between different conformers [77]. The N-myc TAD *R*_2_ values vary across the sequence, ranging between 2.0 s^-1^ and 8.0 s^-1^ with an average of 4.9 s^-1^. Residues with increased *R*_2_ values are Trp50–Phe53, Leu87/Gly89 and Ile111–Glu132, the latter two regions partially align with those shown to exhibit helicity (Fig. 3).

### N-myc peptides for regions of interest

The NMR data highlight sections of N-myc TAD that behave differently to the rest of the protein, be that through transient structure formation and/or altered dynamics. We selected three equal-length 17-residue sections of N-myc for further structural analysis using circular dichroism (CD) spectroscopy (Fig. 5A). Peptides were synthesized for Pro45–Leu61 (MBI); Glu73–Gly89, the region that appears as a helix in the N-myc–Aurora-A crystal structure [31] or ‘Aurora-A-interacting helix’ (AIH) [78]; and Gly119–Gln135, a region with helical propensity that extends beyond MBII

**Figure 5.**
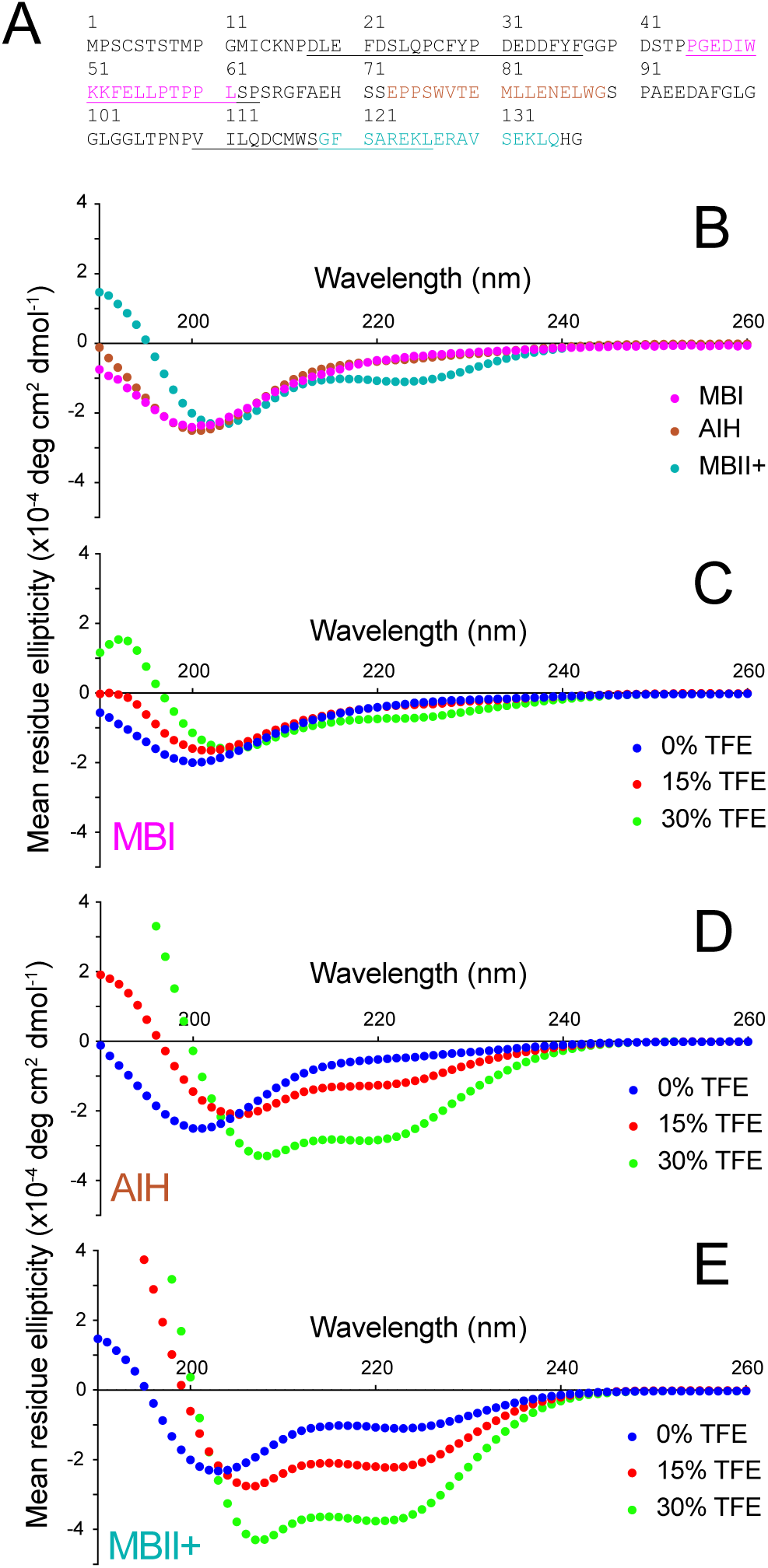
Circular dichroism (CD) spectroscopy of N-myc TAD peptides. (A) N-myc TAD sequence with the positions of the peptides coloured and myc box positions underlined. (B) Overlaid CD spectra for the three peptides in buffer. (C–E) Comparison of CD spectra for the MBI peptide (C), AIH peptide (D) and MBII+ peptide (E) in buffer, 15% TFE and 30% TFE. Experiments were carried out at 5 °C.

(MBII+). At 5 °C, the CD spectra for MBI and AIH appear disordered with a characteristic single minimum in the mean molar residue ellipticity (MRE) at 200 nm [79]. However, MBII+ exhibits a positive band at 190 nm and an increased magnitude negative band at 222 nm (Fig. 5B), characteristics of helicity [80]. An estimate of helical content using the MRE_222_ value shows MBII+ to be 20% helical—which is significant helical content for a short peptide [66]—whereas MBI and AIH might be ∼10% helical. The addition of 2,2,2-fluoroethanol (TFE), a known helix promoter [81], showed that whereas MBI has limited potential to form a helix, AIH can be driven to become very highly helical (estimate >70%), nearly on a par with MBII+ under the same conditions. TFE results can be used to mimic how peptides could interact with other proteins. These results suggest that the AIH region, although not natively helical, can be driven to bind in a helical conformation, as observed in its interaction with Aurora-A, perhaps in a bind-and-fold mechanism. MBII+ exhibits a clear helical signature even without the addition of TFE, marking it as a potential site to bind to partner proteins in this structured form.

MBII is important for interaction of myc proteins with many histone acetyl transferase enzymes, at least in part due to its known role in binding the adaptor protein, transformation/transcription domain-associated protein (TRRAP) [25, 82]. MBII sequences for N-myc (110–126) and c-myc (128–144) have 76% amino acid identity, but beyond that N-myc includes three additional helix promoting residues ‘ERA’ prior to a conserved ‘VSELK’ motif (Fig. S4); the ‘VS’ residues have lower helical propensity and likely act to limit helix propagation [83]. CD spectra for a c-myc MBII+ peptide (residues 137–153, Fig. S4) show it has much weaker helicity (<10% helix) than the equivalent stretch in N-myc. However, as with the AIH in N-myc, the helicity of c-myc MBII+ is significantly enhanced on addition of TFE.

### Interaction with Aurora-A

N-myc TAD is a known binder of the Ser/Thr-kinase Aurora-A [84]. Here, NMR was used to assess the interaction of the full N-myc TAD with Aurora-A. Addition of small volumes of concentrated Aurora-A to a ^15^N-labelled sample of N-myc TAD led to loss in ^1^H–^15^N HSQC spectral quality (Fig. S5). This indicates that, as expected, the two species interact extensively – peak intensity losses in N-myc come about for sites of interaction with Aurora-A due to loss in dynamic freedom (and changes in *R*_2_). From the spectrum of N-myc TAD in complex with Aurora-A, isolated peaks for some residues are completely lost (*e.g.*, Val78, Thr79), strongly indicating them as sites of interaction. However, the initial peak overlap and poor quality of the N-myc–Aurora-A complex spectrum prevented a more thorough analysis. More useful information was gained by running the same Aurora-A NMR titration with ^15^N-labelled truncates of N-myc (Fig. 6). As with the assignment strategy, by dividing the TAD into two sections, the peak overlap is reduced allowing more residues to be individually assessed. It also has the benefit of reducing the overall extent of interaction between Aurora-A and each N-myc truncate and thus the change in *R*_2_ dynamics on binding.

**Figure 6.**
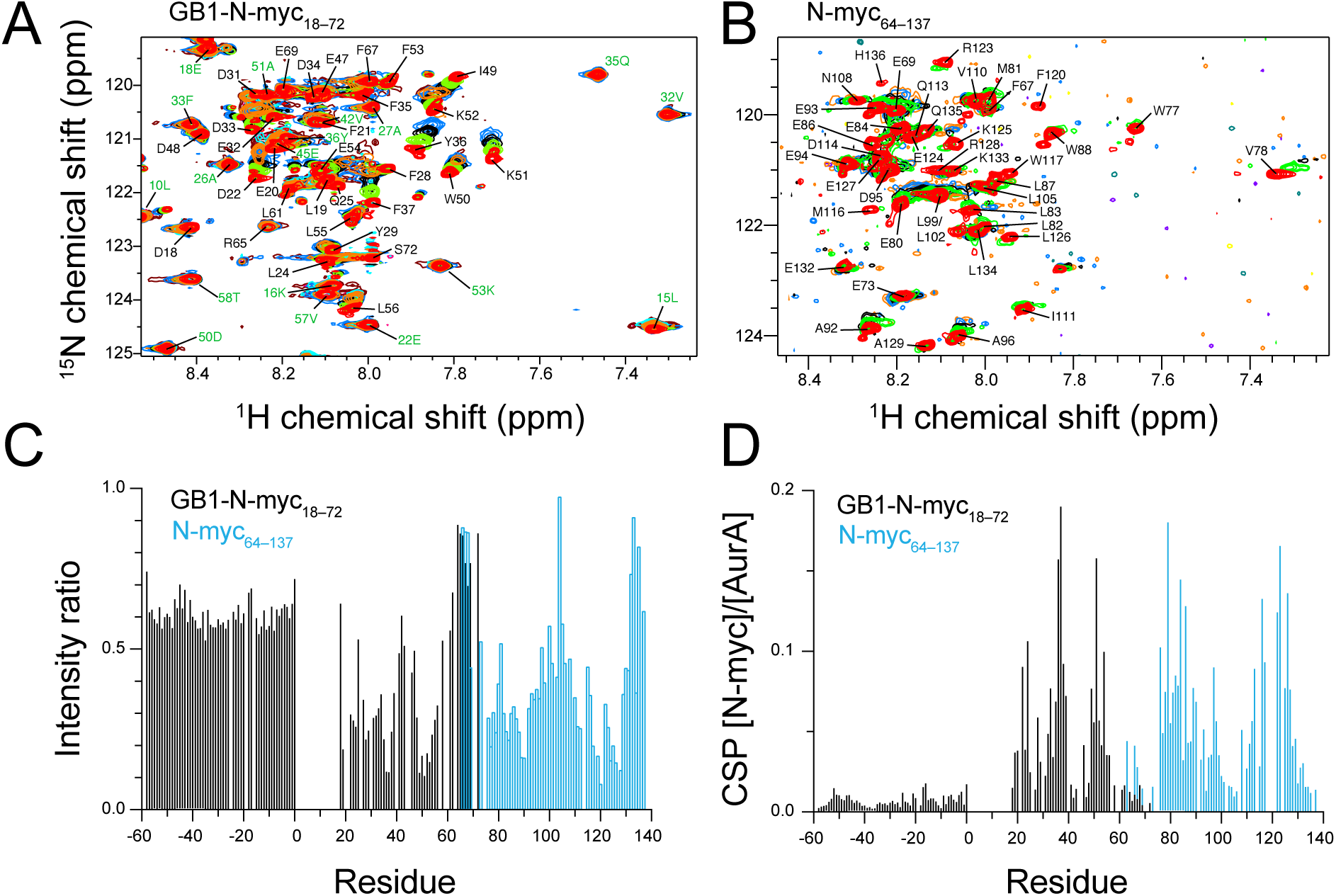
NMR titration experiments for Aurora A kinase domain with N-myc TAD truncates. (A) ^1^H– ^15^N HSQC spectrum of GB1-N-myc_18–72_ (red) and then after addition of increasing amounts of Aurora A kinase domain (lime green, black, sky blue, orange). (B) ^1^H–^15^N HSQC spectrum of N-myc_64–137_ (red) and then after addition of increasing amounts of Aurora A kinase domain (lime green, black, sky blue, orange). (C) Ratio of intensities at a [Aurora A]:[N-myc] molar ratio of 0.2:1 compared to unbound N-myc truncates. The GB1 sequence is renumbered as negative numbers. (D) Changes in chemical shift perturbations as a linear function of [Aurora A]:[N-myc].

A slightly longer ^15^N-labelled N-myc fusion protein (GB1-N-myc_18-72_) was used rather than the assigned GB1-N-myc_18-59_, as this provided some overlap with the C-terminal N-myc_64–137_ construct. Assignment of the C-terminal part of GB1-N-myc_18-72_ was straightforwardly achieved through comparison with full length. On addition of Aurora-A not only peak intensity changes but chemical shift perturbations (CSPs) were observed (Fig. 6A,B). Plotting the ratio of peak intensities for spectra with Aurora-A present to those in initial spectra shows there is a general loss in peak intensity across N-myc samples sequences, but strongly binding regions suffer more significant losses (Fig. 6C). The GB1 peaks provide a useful control since, despite not being involved in the interaction, the overall change in protein tumbling on binding means that their intensities drop to ratios that are consistently ∼0.6 across its sequence. Interestingly, there are regions within the N-myc sequence that maintain significantly higher intensity ratios than in GB1, notably Ser64–Ala68, Gly104 and Glu132–Gln135 at the C-terminus. It is possible that, as seen with c-myc binding to Bin1 [44], these regions are slightly liberated from dynamic intramolecular interactions when neighboring parts of the sequence are occupied in binding to Aurora-A. There appear to be four regions within N-myc with the most significant intensity drops: Leu19–Gly39 (MB0), Asp48–Leu56 (MBI), Ser76–Ser90 (AIH), Val110–Ala129 (MBII). The first three of these regions are known interactors, with Ser76–Gly89 visible in the Aurora-A–N-myc crystal structure [31]. The last region, which aligns with MBII has not been previously characterised, and so the data indicate that the N-myc–Aurora-A interface is more extensive than previously thought. The patterns of CSPs observed on binding mirror those of the intensity drops with non-interacting sites (and GB1) showing small shifts and larger CSP observed for the regions described above (Fig. 6D). The CSP plot shows two peaks within the Leu19–Gly39 region, which could suggest that there are two distinct binding motifs herein separated by a flexible linker.

### In vitro phosphorylation of N-myc TAD

Phosphorylation of the TAD is associated with regulated ubiquitination–proteolysis of N-myc, and NMR provides a means to monitor the individual phosphorylation events catalyzed by different kinases. ERK1 is a proline-directed kinase which becomes stimulated by mitogenic signals and has been shown to specifically phosphorylate Ser62 in c-myc, both *in vivo* and *in vitro* [34, 85]. The ^1^H–^15^N HSQC spectra of N-myc TAD prior to addition and post-incubation with 0.3 μM of ERK1 were compared (Fig. 7A). The ^1^H–^15^N peak for Ser62 undergoes a large chemical shift perturbation (CSP) characteristic of phosphorylation (downfield shift in both ^1^H and ^15^N) [63]. Peaks for residues surrounding this site: Leu61, Ser64, Arg65 and Gly66 also experience significant CSPs, indicating that their chemical environment has changed, due to the presence of the nearby phosphoryl group. Small CSPs were also observed for more distant His residues and their nearest neighbors (His70, Leu71, His136, Gly137). These changes are most likely not associated with ERK1 activity directly but are due to changes in protonation state of the histidines as the pH in the solution subtly decreases due to ATP hydrolysis.

**Figure 7.**
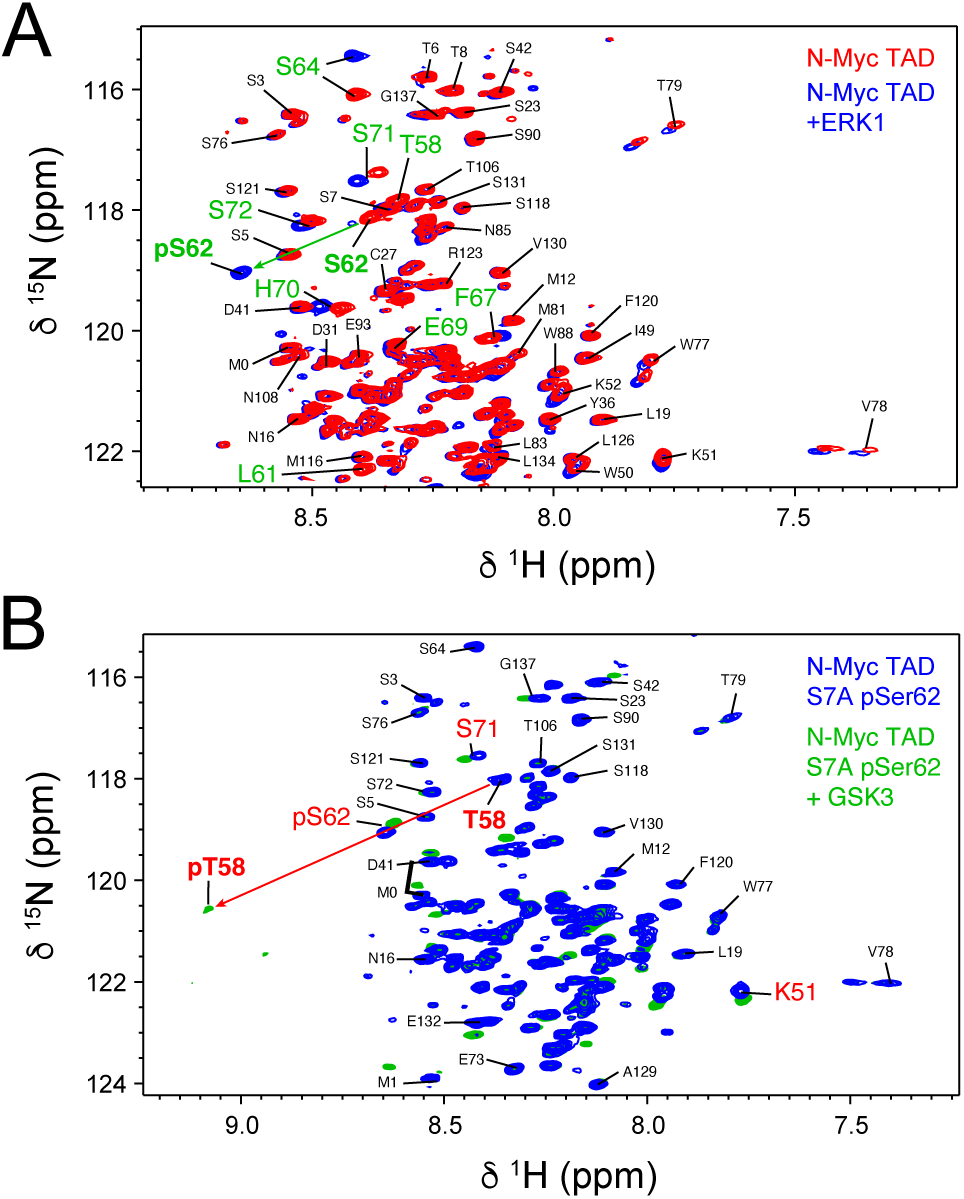
*In vitro* phosphorylation of N-myc TAD followed by NMR. (A) ^1^H–^15^N HSQC spectrum of N-myc TAD (red) and then after addition of ERK1 (blue). (B) ^1^H–^15^N HSQC spectrum of N-myc TAD S7A, pre-phosphorylated by ERK1 (blue) and then after addition of GSK3 (green).

Four additional Ser/Thr residues within N-myc TAD (Thr43, Thr58, Ser90, Thr106) meet the criteria for the ERK1 minimal consensus sequence ([Ser/Thr]–[Pro]) [86]. We therefore examined if these sites were also targeted. Thr43 and Thr106 also undergo phosphorylation by ERK1, albeit on a much slower timescale than Ser62 (Fig. S6). Unlike Ser62, which becomes fully phosphorylated within the time required to collect the spectrum, Thr43 and Thr106 take on the order of 10 and 5 hours, respectively, for their HSQC peaks to diminish to 50% of the original intensity – a timescale unlikely to be physiologically relevant. There was concomitant slow emergence of new, weak, downfield shifted ^1^H–^15^N HSQC peaks associated with these phosphorylated-species. Interestingly, the Ser90 peak did not appreciably change over the course of phosphorylation experiments, behaving in a manner equivalent to other Ser residues that do not meet the minimal consensus sequence (*e.g*. Ser23, Ser118, Ser131). The direct effect of ERK1 on Thr58 was difficult to observe since a small CSP from the phosphorylation of Ser62 causes the ^1^H–^15^N HSQC peak of Thr58 to overlap with that of Ser7. This unfortunate feature also prevented the next step in the myc degradation pathway—phosphorylation of Thr58 by GSK3—from being unambiguously tracked.

To address the peak overlap problem, an N-myc TAD construct incorporating a S7A mutation was produced. The ^1^H–^15^N HSQC spectrum of N-myc TAD^S7A^ overlaps very well with WT N-myc TAD, except for the mutation site itself and peaks for a few surrounding residues. Importantly, the absence of the Ser7 peak meant that Thr58 could be independently tracked throughout. N-myc TAD^S7A^ was subjected to the same initial phosphorylation procedures, phosphorylation with ERK1, followed by addition of 0.3 μM GSK3 monitored using NMR (Fig. 7B). The shifted Thr58 peak after Ser62 phosphorylation did not show obvious loss in peak intensity (Fig. S6B) suggesting that Thr58 in N-myc is not efficiently phosphorylated by ERK1, at least in the presence of a neighboring Ser62/p-Ser62. After addition of GSK3 the Thr58 peak is rapidly and completely lost (Fig. 7B, Fig. S6B), and a new broad peak appears downfield in ^1^H and ^15^N, within the region expected for phospho-species [63]. The action of GSK3 appears to be specific to Thr58. Some neighboring residues (*e.g*., Leu56, Leu61) displayed small CSPs, generally smaller than those observed with the priming phosphorylation of Ser62 (Fig. 7). We confirmed that phosphorylation of Thr58 by GSK3 requires the prior phosphorylation of Ser62 by treating unphosphorylated N-myc TAD with GSK3 and following changes using intact mass spectrometry (Fig. S7). Only low levels of phosphorylated material (+80 Da) were present after ∼3 h. A low level of phosphorylation is consistent with overexpressed GSK3 being able to phosphorylate Thr58 *in vivo*, when a S62A mutation is present [37].

### Doubly phosphorylated N-myc TAD binds the Fbxw7–Skp1 complex

The phospho-degrons of c-myc and N-myc are recognized by Fbxw7–Skp1, which mediates their ubiquitination, targeting them for proteosomal degradation [35, 41]. To establish the importance of individual PTMs on Fbxw7–Skp1 recognition, N-myc TAD p-Ser62 and N-myc TAD p-Ser62 p-Thr58 were generated by *in vitro* phosphorylation using ERK1 and GSK3, the reactions monitored using NMR as described above and stopped by snap freezing in liquid nitrogen. Unphosphorylated N-myc TAD, N-myc TAD p-Ser62, and N-myc TAD p-Thr58 p-Ser62 were incubated with Fbxw7–Skp1 and analytical SEC was employed to ascertain complex formation (Fig. 8). N-myc TAD p-Ser62 did not form a stable complex with Fbw7–Skp1 and the proteins eluted separately at the same elution volumes as individual proteins (Fig. 8A), as confirmed by SDS-PAGE. The interaction between the two species only became apparent when N-myc TAD was di-phosphorylated on Thr58 and Ser62. The N-myc peak almost completely disappears and is found to co-elute with Fbxw7–Skp1 (Fig. 8B). Interestingly, the complex between Fbxw7–Skp1 and N-myc TAD p-Thr58 p-Ser62 elutes later than the Fbxw7–Skp1 complex alone, suggesting that the interaction with N-myc TAD makes the complex more compact despite an increase in its molecular weight.

**Figure 8.**
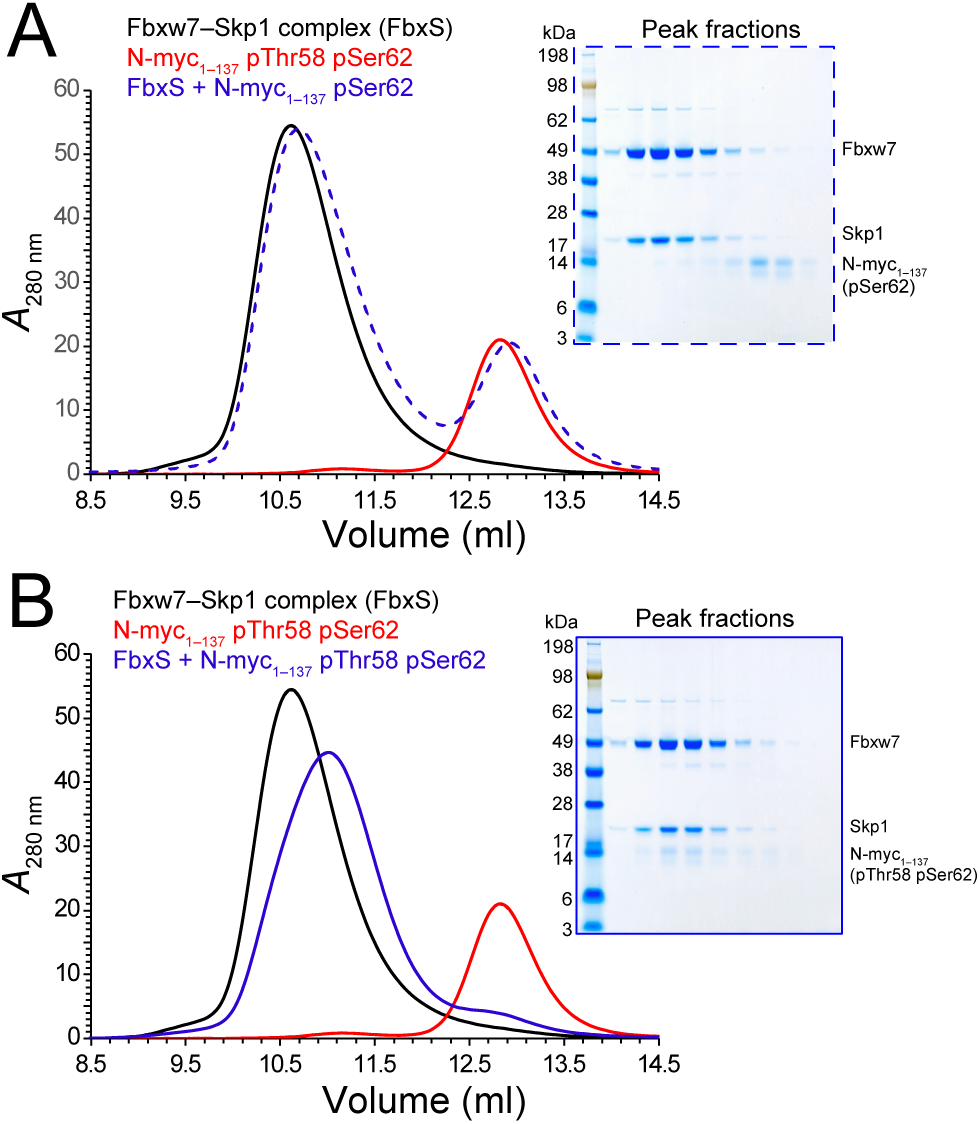
Doubly phosphorylated N-myc TAD (pSer62, pThr58) binds to the Fbxw7–Skp1 complex. Analytical size exclusion chromatography using a Superose12 10/300 GL column was used to assess complex formation between the Fbxw7–Skp1 complex and either singly phosphorylated (pSer62) N-myc TAD (A), or doubly phosphorylated (pSer62, pThr58) N-myc TAD (B). The chromatogram for the complex is shown in blue, while the chromatograms for N-myc alone or Fbxw7–Skp1 alone are shown in red and black, respectively. 0.5 mL fractions were taken at the same positions in both runs and analysed by SDS-PAGE.

## DISCUSSION

Myc transactivation domains are intrinsically disordered regions, vital for the regulation of gene expression, and essential for the oncogenic functions observed in myc proteins. Both N-myc and c-myc interact with a large number of proteins via the TAD to mediate these functions. This region of N-myc also contains the MBI phosphodegron, which is an important determinant of protein stability, and a site which is frequently mutated in N-myc driven cancers [87]. Structural understanding of these regions has been limited to crystal structures of small fragments and NMR studies on an N-terminal portion of c-myc. The assignment of the full TAD of N-myc will be useful to the field for understanding the dynamics and interactions which underpin the oncogenic and physiological functions of this part of N-myc. In this study, we have characterised the dynamics of N-myc in solution, expanded the known interaction region with a key partner and explored the phosphorylation of the MBI phosphodegron.

### Multi-site interaction with Aurora-A perturbs the N-myc TAD

We have previously mapped the interaction between N-myc and Aurora-A using truncated N-myc fragments and determined a crystal structure of N-myc_28–89_ bound to the kinase domain of Aurora-A [31]. Only residues 61–89 were observed in the crystal structure, with residues 76–89 forming an α-helix packed onto the C-lobe of Aurora-A. This region of N-myc matches one of the areas with intrinsic helical propensity shown in the NMR analysis above, suggesting there is a level of ‘templating’, *i.e*., the intrinsic helicity of this region and Aurora-A binding are coupled, although it is difficult to say which feature dominates. The N-terminal section of N-myc was not observed in the crystal structure although FP binding assays showed that N-myc_18–47_ could bind independently to Aurora-A.

NMR studies on this interaction showed the expected shifts in ^1^H–^15^N resonances in the MB0 and the 74–89 helical region, both in terms of intensity and chemical shift. The perturbation of residues close to, or in, MBII was surprising. This may represent a new N-myc–Aurora-A interaction interface which acts in concert with the other two sites. While it is common for myc proteins to bind to their partners using more than one linear motif [30, 53, 88], the use of three motifs spanning across ∼120 residues is unusual.

### Two-site, specific phosphorylation of N-myc MBI

Using in-NMR kinase assays, we established the strong preference of ERK1 for Ser62 and the dependence of GSK3 on this priming event for efficient and selective phosphorylation of Thr58. Phosphorylation of Ser62 and subsequently double phosphorylation of Ser62 and Thr58 allowed for unambiguous assignment of the ^1^H–^15^N resonances of the phosphorylated residues. The absence of chemical shift perturbations other than in MBI is strongly suggestive that there are not any major changes in N-myc TAD structure driven by either phosphorylation event. This was not entirely unexpected, given that phosphorylation at Ser62 in c-myc 1–88 had no major effects on the chemical shifts outside of MBI either. That this doubly phosphorylated material forms a stable complex with Fbxw7–Skp1 corroborates the findings of Welcker and colleagues [37] who found that doubly phosphorylated MBI was the predominant c-myc species found bound to Fbxw7 in cells and additionally determined the structure of a doubly phosphorylated MBI peptide in complex with Fbxw7–Skp1.

The N-myc TAD is thought to be post-translationally modified on at least three other sites. Two of these sites are close to the MBI phosphodegron; S64 which is phosphorylated and R65 which is thought to be methylated [89, 90]. A site close to MBII, K133, is also thought to be acetylated [91]. The approach taken to monitor phosphodegron phosphorylation here can also be used to assess post translational modifications on these other sites, either to determine their potential roles in changing phosphorylation dynamics of MBI, in changing N-myc TAD dynamics, or in interacting with binding partners.

### Ordered regions of N-myc TAD beyond the myc boxes participate in protein–protein interactions

Analysis of the solution dynamics of N-myc TAD indicate that is not a fully disordered polypeptide: in several regions of the TAD, its motions are less dynamic on the ps–ns timescale than would be expected for a fully disordered polypeptide, resulting in heterogeneity in peak intensities across the TAD in NMR experiments. Regions of N-myc 1–137 involved in protein–protein interactions (notably AIH and MBII) have clear helical propensity, and the protein samples these secondary structure conformations in solution. We do not yet understand the consequences of this conformational sampling, however for c-myc it has been observed that binding of Bin1 to one site increased flexibility elsewhere in the protein [44]; similar behaviour was observed here when N-myc binds Aurora-A. Molecular dynamics simulations of both c-myc 1–88 and the full-length c-myc–Max complex also suggest that myc can sample many conformations and that conformational sampling in their transactivation domain may be a viable target for drugging c-myc [92]. It remains to be determined if the constrained dynamics of myc proteins have roles in myc biology. However, this feature of constrained dynamics at the ps–ns timescale is not observed in all intrinsically disordered TADs. A well-studied example in NMR is that of the human p53 TAD. This appears to be highly dynamic, as demonstrated by its hetNOE values which are at, or below, zero [93, 94].

Chemical shift data indicate the presence of helical propensity in residues 77–86 inclusive and 122–132 inclusive. They lie in a region where the equivalent sequence in c-myc has yet to be fully structurally studied [33] (Fig. S8A). We therefore compared the secondary structure characteristics of this region in the two proteins, starting with AlphaFold2 predictions (Fig. S8B–D). The crystal structure of N-myc bound to Aurora-A kinase demonstrated that residues 75–89 bind to the kinase in an α-helical conformation. While the chemical shift data indicate that this region (the AIH) is sampling helical conformations in solution, CD data for the AIH peptide indicate that there are very low levels of helicity in solution prior to the addition of TFE. This suggests that significant folding upon binding may be required for this region to interact as a helix. The AlphaFold2 model for N-myc (https://alphafold.ebi.ac.uk/entry/P04198) predicts an α-helix but with ‘low’ confidence (average pLDDT score 58.2) from 76–88 (Fig. S8B & D). It has been shown that AlphaFold2 is able to identify conditional folding, particularly of helices, in IDRs [95], *i.e*., formally disordered regions within proteins that are able to fold under certain conditions, typically when bound to another protein chain. Examples of this can be seen in the same myc Alphafold2 models: the C-termini appear as helix-loop-helix structures that would only occur if the binding partner, Max, were present. AGADIR [96] (http://agadir.crg.es) predicts that the AIH region of N-myc will have low helicity in isolation (Fig. S8E). The interaction with Aurora-A can be reinforced using helix-constrained N-myc AIH peptides [78], suggesting that a level of pre-organization can facilitate binding. Although there is limited sequence identity between N-myc and c-myc in the region after MBI, c-myc, in a similar position to the N-myc AIH, has been crystallised interacting as a helix with the TBP–TAF1 complex (c-myc 97–107) [30]. The helices do share a ΦVTEΦL sequence pattern (Fig. S8A), but c-myc lacks the two Trp residues (Trp77 and Trp88) that are important for the N-myc–Aurora-A interaction [31]. Thus, although this region of the TAD is not within a myc box, it has some conserved sequence and structural features which may point to a conserved but unknown interaction. Furthermore, the sequence differences allow for diversification of interactions specialised to each myc paralog; different partners can bind to equivalent sites in different myc proteins (*e.g*., TBP–TAF1 for c-myc and Aurora-A for N-myc).

AlphaFold2 predicts a helix for residues 122–137 in N-myc with high confidence (average pLDDT score 71.2, Fig. S8B & D). Likewise the c-myc AlphaFold2 model (https://alphafold.ebi.ac.uk/entry/P01106) predicts a helix at the same ‘MBII+’ position as in N-myc (Fig. S8D), albeit with reduced confidence (average pLDDT score 64.0). The Alphafold2 model shows a small helix just prior to MBII in c-myc but not in N-myc, presumably being disfavoured due to a pair of Pro residues. The first five residues of MBII+ are in the highly conserved MBII region of the protein, with approximately 3 helical turns beyond the MBII region. CD demonstrated that the N-myc MBII+ peptide has significant helical content in solution even in the absence of TFE, suggesting that it may not need significant folding-upon-binding to interact as a helix. In contrast, c-myc MBII+ has negligible helicity in the absence of TFE. In line with these finding, AGADIR predicts N-myc to be much more helical in this region than c-myc (Fig. S8E). It is possible that the helix, where present, acts as a single entity upon interaction with binding partners. We could therefore consider the functional unit of MBII to extend into this helix, comprising 110–137 in N-myc but to varying degrees in other myc homologues.

Overall, our approach uncovered specific structural features and dynamics of the N-myc TAD that mediate protein–protein interactions, which are conserved to varying degrees in c-myc. Notably, the most striking features involve residues outside the canonical myc boxes, highlighting the importance of structural context in understanding the conserved and divergent functions of myc proteins.

## Data availability

Chemical shift data for GB1-N-myc_18–59_, N-myc TAD, and N-myc_64–137_ were deposited in the Biological Magnetic Resonance Data Bank (BMRB) entries 52047, 52066 and 52067, respectively.

## Supporting information

This article contains supporting information.

## Conflict of interest

The authors declare that they have no conflict of interest with the contents of this article.

## Acknowledgements

We would like to acknowledge G. Nasir Khan for maintenance and assistance with the circular dichroism spectrometer.

## Author contributions

E.R., E.L., M.S.A. and M.W.R. provided proteins and reagents, E.R., E.L. and M.B. collected data and analysed results, A.P.K. assisted with NMR data collection. R.B. conceived the project. All authors contributed to the writing and editing of the manuscript.

## Funding

MRC DiMeN programme for DTP studentships (to E.R.). CRUK Programme Award (C24461/A23302 to R.B.). BBSRC sLoLa (BB/V003577/1 to R.B.). MRC Project Award (MR/V029975/1 to R.B. and E.L.). Facilities for NMR spectroscopy and CD spectroscopy were funded by the University of Leeds (ABSL award) and Wellcome Trust (108466/Z/15/Z, 094232/Z/10/Z).

## Notes

https://bmrb.io/data_library/summary/index.php?bmrbId=52047

https://bmrb.io/data_library/summary/index.php?bmrbId=52066

https://bmrb.io/data_library/summary/index.php?bmrbId=52067

### Competing Interest Statement

The authors have declared no competing interest.

## REFERENCES

1. Hurlin, P.J., Control of vertebrate development by MYC. Cold Spring Harbor perspectives in medicine, 2013. 3(9): p. a014332–a014332.

2. Casey, S.C., V. Baylot, and D.W. Felsher, The MYC oncogene is a global regulator of the immune response. Blood, 2018. 131(18): p. 2007–2015.

3. Dang, C.V., MYC, metabolism, cell growth, and tumorigenesis. Cold Spring Harbor perspectives in medicine, 2013. 3(8): p. a014217.

4. Schwab, M., et al., Amplified DNA with limited homology to myc cellular oncogene is shared by human neuroblastoma cell lines and a neuroblastoma tumour. Nature, 1983. 305(5931): p. 245–248.

5. Rickman, D.S., J.H. Schulte, and M. Eilers, The Expanding World of N-MYC–Driven Tumors. Cancer Discovery, 2018. 8(2): p. 150–163.

6. Brodeur, G.M., et al., Amplification of N-myc in Untreated Human Neuroblastomas Correlates with Advanced Disease Stage. Science, 1984. 224(4653): p. 1121–1124.

7. Schwab, M., Amplification of N-myc as a prognostic marker for patients with neuroblastoma. Seminars in cancer biology, 1993. 4(1): p. 13–18.

8. Wang, L.L., et al., Augmented expression of MYC and/or MYCN protein defines highly aggressive MYC-driven neuroblastoma: a Children’s Oncology Group study. British journal of cancer, 2015. 113(1): p. 57–63.

9. Nie, Z., et al., *c-Myc is a universal amplifier of expressed genes in lymphocytes and embryonic stem cells*. Cell, 2012. 151(1): p. 68–79.

10. Nie, Z., et al., Dissecting transcriptional amplification by MYC. eLife, 2020. 9: p. e52483.

11. Lin, C.Y., et al., Transcriptional amplification in tumor cells with elevated c-Myc. Cell, 2012. 151(1): p. 56–67.

12. Gomez-Roman, N., et al., Direct activation of RNA polymerase III transcription by c-Myc. Nature, 2003. 421(6920): p. 290–294.

13. Grandori, C., et al., *c-Myc binds to human ribosomal DNA and stimulates transcription of rRNA genes by RNA polymerase I*. Nature Cell Biology, 2005. 7(3): p. 311–318.

14. Arabi, A., et al., *c-Myc associates with ribosomal DNA and activates RNA polymerase I transcription*. Nature Cell Biology, 2005. 7(3): p. 303–310.

15. de Pretis, S., et al., Integrative analysis of RNA polymerase II and transcriptional dynamics upon MYC activation. Genome research, 2017. 27(10): p. 1658–1664.

16. Herkert, B. and M. Eilers, Transcriptional repression: the dark side of myc. Genes & cancer, 2010. 1(6): p. 580–586.

17. Staller, P., et al., Repression of p15INK4b expression by Myc through association with Miz-1. Nature Cell Biology, 2001. 3(4): p. 392–399.

18. Prendergast, G.C., D. Lawe, and E.B. Ziff, Association of Myn, the murine homolog of Max, with c-Myc stimulates methylation-sensitive DNA binding and ras cotransformation. Cell, 1991. 65(3): p. 395–407.

19. Blackwood, E.M. and R.N. Eisenman, Max: A Helix-Loop-Helix Zipper Protein That Forms a Sequence-Specific DNA-Binding Complex with Myc. Science, 1991. 251(4998): p. 1211–1217.

20. Kerkhoff, E., K. Bister, and K.H. Klempnauer, Sequence-specific DNA binding by Myc proteins. Proceedings of the National Academy of Sciences of the United States of America, 1991. 88(10): p. 4323–4327.

21. Prendergast, G.C. and E.B. Ziff, Methylation-Sensitive Sequence-Specific DNA Binding by the c-Myc Basic Region. Science, 1991. 251(4990): p. 186–189.

22. Blackwell, T.K., et al., Sequence-Specific DNA Binding by the c-Myc Protein. Science, 1990. 250(4984): p. 1149–1151.

23. Nair, S.K. and S.K. Burley, X-Ray Structures of Myc-Max and Mad-Max Recognizing DNA. Cell, 2003. 112(2): p. 193–205.

24. Büchel, G., et al., Association with Aurora-A Controls N-MYC-Dependent Promoter Escape and Pause Release of RNA Polymerase II during the Cell Cycle. Cell reports, 2017. 21(12): p. 3483–3497.

25. Kalkat, M., et al., MYC Protein Interactome Profiling Reveals Functionally Distinct Regions that Cooperate to Drive Tumorigenesis. Molecular Cell, 2018. 72(5): p. 836–848.e7.

26. Malynn, B.A., et al., N-myc can functionally replace c-myc in murine development, cellular growth, and differentiation. Genes & Development, 2000. 14(11): p. 1390–1399.

27. Kato, G.J., et al., An amino-terminal c-myc domain required for neoplastic transformation activates transcription. Molecular and cellular biology, 1990. 10(11): p. 5914–5920.

28. Spotts, G.D., et al., Identification of downstream-initiated c-Myc proteins which are dominant-negative inhibitors of transactivation by full-length c-Myc proteins. Molecular and cellular biology, 1997. 17(3): p. 1459–1468.

29. Junjiao, Y., et al., Phase separation of Myc differentially regulates gene transcription. bioRxiv, 2022: p. 2022.06.28.498043.

30. Wei, Y., et al., Multiple direct interactions of TBP with the MYC oncoprotein. Nature Structural & Molecular Biology, 2019. 26(11): p. 1035–1043.

31. Richards, M.W., et al., Structural basis of N-Myc binding by Aurora-A and its destabilization by kinase inhibitors. Proceedings of the National Academy of Sciences of the United States of America, 2016. 113(48): p. 13726–13731.

32. Roeschert, I., et al., Combined inhibition of Aurora-A and ATR kinase results in regression of MYCN-amplified neuroblastoma. Nature cancer, 2021. 2(3): p. 312–326.

33. Schutz, S., et al., Intrinsically Disordered Regions in the Transcription Factor MYC:MAX Modulate DNA Binding via Intramolecular Interactions. Biochemistry, 2024.

34. Sears, R., et al., Multiple Ras-dependent phosphorylation pathways regulate Myc protein stability. Genes & development, 2000. 14(19): p. 2501–2514.

35. Sjostrom, S.K., et al., The Cdk1 Complex Plays a Prime Role in Regulating N-Myc Phosphorylation and Turnover in Neural Precursors. Developmental Cell, 2005. 9(3): p. 327–338.

36. Hydbring, P., et al., Phosphorylation by Cdk2 is required for Myc to repress Ras-induced senescence in cotransformation. Proceedings of the National Academy of Sciences of the United States of America, 2010. 107(1): p. 58–63.

37. Welcker, M., et al., Two diphosphorylated degrons control c-Myc degradation by the Fbw7 tumor suppressor. Science advances, 2022. 8(4): p. eabl7872–eabl7872.

38. Ciechanover, A., et al., Degradation of MYCN oncoprotein by the ubiquitin system. Progress in clinical and biological research, 1991. 366: p. 37–43.

39. Flinn, E.M., C.M. Busch, and A.P. Wright, myc boxes, which are conserved in myc family proteins, are signals for protein degradation via the proteasome. Molecular and cellular biology, 1998. 18(10): p. 5961–5969.

40. Gross-Mesilaty, S., et al., Basal and human papillomavirus E6 oncoprotein-induced degradation of Myc proteins by the ubiquitin pathway. Proceedings of the National Academy of Sciences of the United States of America, 1998. 95(14): p. 8058–8063.

41. Yada, M., et al., Phosphorylation-dependent degradation of c-Myc is mediated by the F-box protein Fbw7. The EMBO journal, 2004. 23(10): p. 2116–2125.

42. Hann, S.R. and R.N. Eisenman, Proteins encoded by the human c-myc oncogene: differential expression in neoplastic cells. Molecular and cellular biology, 1984. 4(11): p. 2486–2497.

43. Wei, Y., et al., The MYC oncoprotein directly interacts with its chromatin cofactor PNUTS to recruit PP1 phosphatase. Nucleic acids research, 2022. 50(6): p. 3505–3522.

44. Andresen, C., et al., Transient structure and dynamics in the disordered c-Myc transactivation domain affect Bin1 binding. Nucleic acids research, 2012. 40(13): p. 6353–6366.

45. Macek, P., et al., Myc phosphorylation in its basic helix-loop-helix region destabilizes transient α-helical structures, disrupting Max and DNA binding. The Journal of biological chemistry, 2018. 293(24): p. 9301–9310.

46. Panova, S., et al., Mapping Hidden Residual Structure within the Myc bHLH-LZ Domain Using Chemical Denaturant Titration. Structure, 2019. 27(10): p. 1537–1546.e4.

47. Lavigne, P., et al., Insights into the mechanism of heterodimerization from the 1H-NMR solution structure of the c-Myc-Max heterodimeric leucine zipper. Journal of Molecular Biology, 1998. 281(1): p. 165–181.

48. Kızılsavaş, G., et al., *¹H, ¹³C,* and ¹⁵N backbone and side chain resonance assignments of the C-terminal DNA binding and dimerization domain of v-Myc. Biomolecular NMR assignments, 2013. 7(2): p. 321–324.

49. Baminger, B., et al., Letter to the editor: Backbone assignment of the dimerization and DNA-binding domain of the oncogenic transcription factor v-Myc in complex with its authentic binding partner Max. Journal of Biomolecular NMR, 2004. 30(3): p. 361–362.

50. Sammak, S., et al., The structure of INI1/hSNF5 RPT1 and its interactions with the c-MYC:MAX heterodimer provide insights into the interplay between MYC and the SWI/SNF chromatin remodeling complex. The FEBS journal, 2018. 285(22): p. 4165–4180.

51. Sammak, S., et al., Crystal Structures and Nuclear Magnetic Resonance Studies of the Apo Form of the c-MYC:MAX bHLHZip Complex Reveal a Helical Basic Region in the Absence of DNA. Biochemistry, 2019. 58(29): p. 3144–3154.

52. Schütz, S., et al., The Disordered MAX N-terminus Modulates DNA Binding of the Transcription Factor MYC:MAX. Journal of Molecular Biology, 2022. 434(22): p. 167833.

53. Helander, S., et al., Pre-Anchoring of Pin1 to Unphosphorylated c-Myc in a Fuzzy Complex Regulates c-Myc Activity. Structure (London, England : 1993), 2015. 23(12): p. 2267–2279.

54. Bagneris, C., et al., Crystal structure of a vFlip-IKKgamma complex: insights into viral activation of the IKK signalosome. Mol Cell, 2008. 30(5): p. 620–31.

55. Huth, J.R., et al., Design of an expression system for detecting folded protein domains and mapping macromolecular interactions by NMR. Protein Sci, 1997. 6(11): p. 2359–64.

56. Burgess, S.G. and R. Bayliss, The structure of C290A:C393A Aurora A provides structural insights into kinase regulation. Acta Crystallogr F Struct Biol Commun, 2015. 71(Pt 3): p. 315–9.

57. Delaglio, F., et al., NMRPipe: A multidimensional spectral processing system based on UNIX pipes. Journal of Biomolecular NMR, 1995. 6(3): p. 277–293.

58. Vranken, W.F., et al., *The CCPN data model for NMR spectroscopy: Development of a software pipeline.* Proteins: Structure, Function, and Bioinformatics, 2005. 59(4): p. 687–696.

59. Hendus-Altenburger, R., et al., Random coil chemical shifts for serine, threonine and tyrosine phosphorylation over a broad pH range. J Biomol NMR, 2019. 73(12): p. 713–725.

60. Shen, Y. and A. Bax, Protein backbone and sidechain torsion angles predicted from NMR chemical shifts using artificial neural networks. J Biomol NMR, 2013. 56(3): p. 227–41.

61. Ahlner, A., et al., PINT: a software for integration of peak volumes and extraction of relaxation rates. Journal of Biomolecular NMR, 2013. 56(3): p. 191–202.

62. Niklasson, M., et al., Comprehensive analysis of NMR data using advanced line shape fitting. Journal of Biomolecular NMR, 2017. 69(2): p. 93–99.

63. Selenko, P., et al., In situ observation of protein phosphorylation by high-resolution NMR spectroscopy. Nat Struct Mol Biol, 2008. 15(3): p. 321–9.

64. Batchelor, M., et al., *alpha-Helix stabilization by co-operative side chain charge-reinforced interactions to phosphoserine in a basic kinase-substrate motif*. Biochem J, 2022. 479(5): p. 687–700.

65. Errington, N. and A.J. Doig, A phosphoserine-lysine salt bridge within an alpha-helical peptide, the strongest alpha-helix side-chain interaction measured to date. Biochemistry, 2005. 44(20): p. 7553–8.

66. Baker, E.G., et al., Local and macroscopic electrostatic interactions in single alpha-helices. Nat Chem Biol, 2015. 11(3): p. 221–8.

67. Dyson, H.J. and P.E. Wright, Intrinsically unstructured proteins and their functions. Nat Rev Mol Cell Biol, 2005. 6(3): p. 197–208.

68. Carugo, O., Amino acid composition and protein dimension. Protein Sci, 2008. 17(12): p. 2187–91.

69. Sigler, P.B., Transcriptional activation. Acid blobs and negative noodles. Nature, 1988. 333(6170): p. 210–2.

70. Gibbs, E.B., E.C. Cook, and S.A. Showalter, Application of NMR to studies of intrinsically disordered proteins. Archives of Biochemistry and Biophysics, 2017. 628: p. 57–70.

71. Foster, M.P., C.A. McElroy, and C.D. Amero, Solution NMR of large molecules and assemblies. Biochemistry, 2007. 46(2): p. 331–340.

72. Takeuchi, K., et al., Nitrogen-detected CAN and CON experiments as alternative experiments for main chain NMR resonance assignments. Journal of Biomolecular NMR, 2010. 47(4): p. 271–282.

73. Wishart, D.S., B.D. Sykes, and F.M. Richards, The chemical shift index: a fast and simple method for the assignment of protein secondary structure through NMR spectroscopy. Biochemistry, 1992. 31(6): p. 1647–1651.

74. Kay, L.E., D.A. Torchia, and A. Bax, Backbone dynamics of proteins as studied by nitrogen-15 inverse detected heteronuclear NMR spectroscopy: application to staphylococcal nuclease. Biochemistry, 1989. 28(23): p. 8972–8979.

75. Yao, J., et al., NMR Structural and Dynamic Characterization of the Acid-Unfolded State of Apomyoglobin Provides Insights into the Early Events in Protein Folding. Biochemistry, 2001. 40(12): p. 3561–3571.

76. Estrada, D.F., et al., The Structure of the Hantavirus Zinc Finger Domain is Conserved and Represents the Only Natively Folded Region of the Gn Cytoplasmic Tail. Front Microbiol, 2011. 2: p. 251.

77. Kovermann, M., P. Rogne, and M. Wolf-Watz, Protein dynamics and function from solution state NMR spectroscopy. Q Rev Biophys, 2016. 49: p. e6.

78. Dawber, R.S., et al., Inhibition of Aurora-A/N-Myc Protein-Protein Interaction Using Peptidomimetics: Understanding the Role of Peptide Cyclization. Chembiochem, 2024. 25(2): p. e202300649.

79. Greenfield, N.J., Using circular dichroism spectra to estimate protein secondary structure. Nature Protocols, 2006. 1(6): p. 2876–2890.

80. Woody, R.W., Circular dichroism, in Methods in Enzymology. 1995, Academic Press. p. 34–71.

81. Cammers-Goodwin, A., et al., Mechanism of Stabilization of Helical Conformations of Polypeptides by Water Containing Trifluoroethanol. Journal of the American Chemical Society, 1996. 118(13): p. 3082–3090.

82. McMahon, S.B., et al., The novel ATM-related protein TRRAP is an essential cofactor for the c-Myc and E2F oncoproteins. Cell, 1998. 94(3): p. 363–74.

83. Chakrabartty, A., T. Kortemme, and R.L. Baldwin, Helix propensities of the amino acids measured in alanine-based peptides without helix-stabilizing side-chain interactions. Protein Sci, 1994. 3(5): p. 843–52.

84. Otto, T., et al., Stabilization of N-Myc is a critical function of Aurora A in human neuroblastoma. Cancer Cell, 2009. 15(1): p. 67–78.

85. Yeh, E., et al., A signalling pathway controlling c-Myc degradation that impacts oncogenic transformation of human cells. Nat Cell Biol, 2004. 6(4): p. 308–18.

86. Gonzalez, F.A., D.L. Raden, and R.J. Davis, Identification of substrate recognition determinants for human ERK1 and ERK2 protein kinases. J Biol Chem, 1991. 266(33): p. 22159–63.

87. Bonilla, X., et al., Genomic analysis identifies new drivers and progression pathways in skin basal cell carcinoma. Nat Genet, 2016. 48(4): p. 398–406.

88. Zhang, N., et al., MYC interacts with the human STAGA coactivator complex via multivalent contacts with the GCN5 and TRRAP subunits. Biochim Biophys Acta, 2014. 1839(5): p. 395–405.

89. Eberhardt, A., et al., Protein arginine methyltransferase 1 is a novel regulator of MYCN in neuroblastoma. Oncotarget, 2016. 7(39): p. 63629–63639.

90. Zhou, H., et al., Toward a comprehensive characterization of a human cancer cell phosphoproteome. J Proteome Res, 2013. 12(1): p. 260–71.

91. Mertins, P., et al., Integrated proteomic analysis of post-translational modifications by serial enrichment. Nat Methods, 2013. 10(7): p. 634–7.

92. Lama, D., et al., A druggable conformational switch in the c-MYC transactivation domain. Nat Commun, 2024. 15(1): p. 1865.

93. Lee, H., et al., Local structural elements in the mostly unstructured transcriptional activation domain of human p53. J Biol Chem, 2000. 275(38): p. 29426–32.

94. Schrag, L.G., et al., Cancer-Associated Mutations Perturb the Disordered Ensemble and Interactions of the Intrinsically Disordered p53 Transactivation Domain. J Mol Biol, 2021. 433(15): p. 167048.

95. Alderson, T.R., et al., Systematic identification of conditionally folded intrinsically disordered regions by AlphaFold2. Proc Natl Acad Sci U S A, 2023. 120(44): p. e2304302120.

96. Munoz, V. and L. Serrano, Elucidating the folding problem of helical peptides using empirical parameters. Nat Struct Biol, 1994. 1(6): p. 399–409.

