## Supplemental Information for "Exploring the dynamics and interactions of the N-myc transactivation domain through solution NMR"

**SUPPLEMENTARY INFORMATION**

**Table S1.** List of NMR experiments for each sample with temperature and spectrometer used.

| Spectrum | N-myc_1–137_ (TAD) | | GB1-N-myc_18–59_ | | N-myc_64–137_ | | GB1-N-myc_18–72_ | |
| --- | --- | --- | --- | --- | --- | --- | --- | --- |
|  | Temp. (°C) | Spec. | Temp. (°C) | Spec. | Temp. (°C) | Spec. | Temp. (°C) | Spec. |
| ^1^H-^15^N-HSQC | 10–37 | 750 600 950 | 15 | 600 | 10 | 750 | 15, 37 | 600 |
| HNCO | 10 | 750 | 15 | 600 | 10 | 750^*^ |  |  |
| HNcaCO | 10 | 750 | 15 | 600 | 10 | 750^*^ |  |  |
| HNcoCA | 10 | 750 | 15 | 600 | 10 | 750^*^ |  |  |
| HNCA | 10 | 750 | 15 | 600 | 10 | 750^*^ |  |  |
| HNcocaCB | 10 | 750 | 15 | 600 | 10 | 750^*^ |  |  |
| HNcaCB | 10 | 750 | 15 | 600 | 10 | 750^*^ |  |  |
| ^1^H-^13^C-HSQC | 10 | 750 | 15 | 600 |  |  |  |  |
| HBHAcoNH | 10 | 750 | 15 | 600 |  |  |  |  |
| CON (C-det.) | 10 | 950 |  |  |  |  |  |  |
| hCACO (C-det.) | 10 | 950 |  |  |  |  |  |  |
| CAnCO (C-det) | 10 | 950 |  |  |  |  |  |  |
| CAN (N-det.) | 10 | 950 |  |  |  |  |  |  |
| ^15^N T1 | 10 | 600 |  |  |  |  |  |  |
| ^15^N T2 | 10 | 600 |  |  |  |  |  |  |
| ^15^N–[^1^H]-hetNOE | 10 | 600 |  |  |  |  |  |  |
| ^1^H–^15^N-HSQC Aurora A titration | 25 | 750 |  |  | 35 | 750 | 37 | 750 |
| ^1^H–^15^N-HSQC ERK1, GSK3 phosphorylation | 10 | 600 |  |  |  |  |  |  |
| ^*^ Recorded using 25% non-uniform sampling (NUS). Reconstructed using NMRpipe. | | | | | | | | |

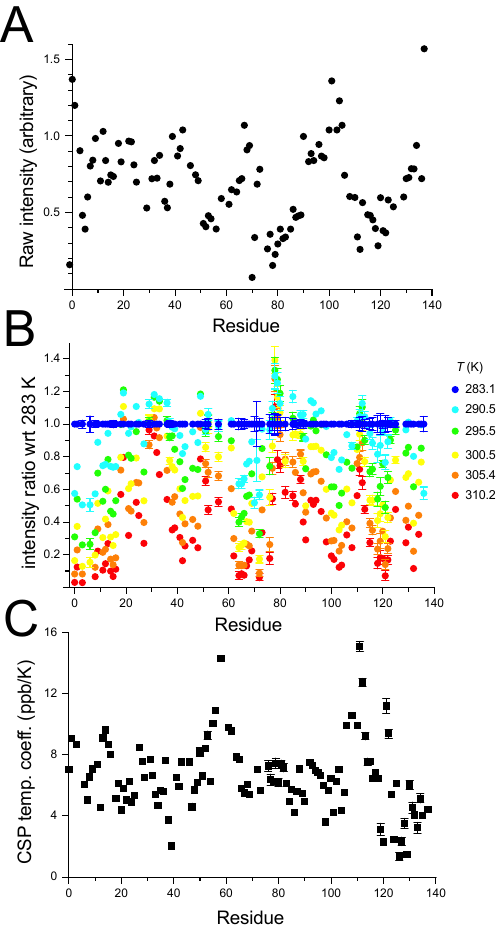

**Figure S1.** Meta-analysis from ^1^H–^15^N HSQC spectra of N-Myc TAD. (A) Raw intensity measurements for peaks across the sequence. Comparatively low intensity (broad) peaks were observed for Lys51–Leu56, Ser76–Gly89 and Ile111–Ala122 regions. (B) Peak intensities as a function of temperature, plotted as a ratio to the peak intensity at 10 °C. (C) Temperature coefficients for HSQC peaks. The plot shows the change in peak position (or chemical shift perturbation, CSP) as a function of temperature.

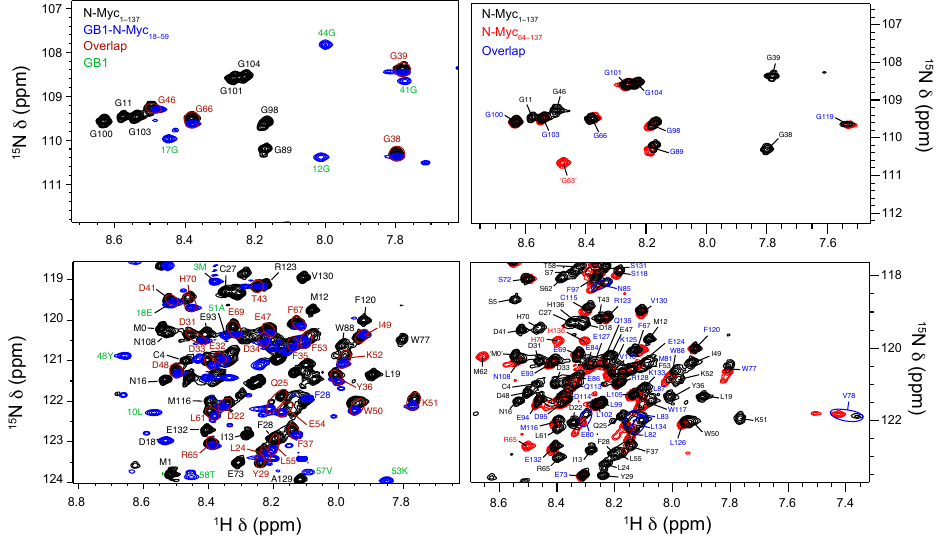

**Figure S2.** A comparison ^1^H–^15^N HSQC spectra of N-myc TAD and the two truncated variants GB1-N-myc_18–59_ (left) and N-myc_64–137_ (right). The upper plots show the glycine region of the spectrum and the lower plots show part of the central region. For the GB1-N-Myc_18–59_ comparison, peaks from the GB1 part of the sequence are labelled in green; overlapping peaks within the 18–59 sequence are labelled in red. For the N-myc_64–137_ comparison, overlapping peaks within the 64–137 sequence are labelled in blue.

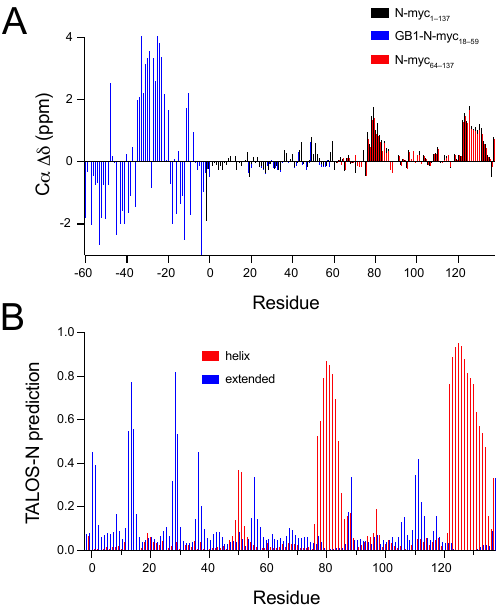

**Figure S3.** (A) A comparison of the Cα secondary shifts for N-myc TAD (black) and the truncated variants GB1-N-myc_18–59_ (blue) N-myc_64–137_ (red). The overlap is excellent in both cases indicating that the structural propensities are likely to the same in the truncated versions. Note particularly the matching of the weak extended propensities between residues 20–35 for GB1-N-myc_18–59_ and the matching helical propensity regions for N-myc_64–137_. It is also worthwhile comparing the secondary shifts observed in N-myc with the fully-folded helix and β-strands in GB1; for instance, the GB1 helix has Δδ values 3–4 times those in the regions with helical propensity in N-myc. (B) TALOS-N prediction of secondary structure with blue bars corresponding to residues with extended/strand propensities and red bars corresponding to those with helicity.

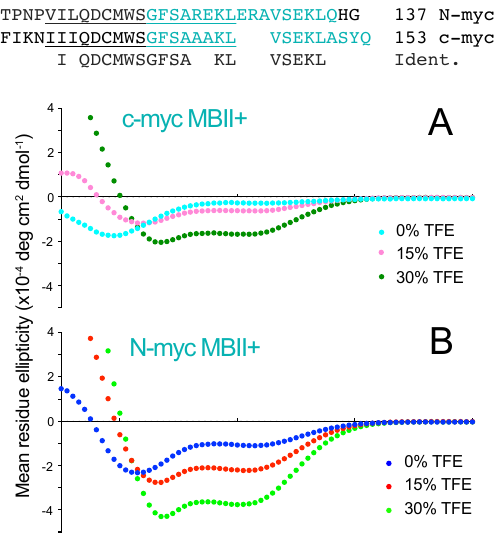

**Figure S4.** Circular dichroism (CD) spectroscopy of c-myc MBII+ peptide. Sequence alignment for the MBII (underlined) and MBII+ regions (coloured teal) in N-myc and c-myc. (A) Comparison of CD spectra for the c-myc MBII+ peptide in buffer, 15% TFE and 30% TFE. (B) To facilitate comparison, spectra for the N-myc MBII+ (already shown in Fig. 5E) are repeated here. Experiments were carried out at 5 °C.

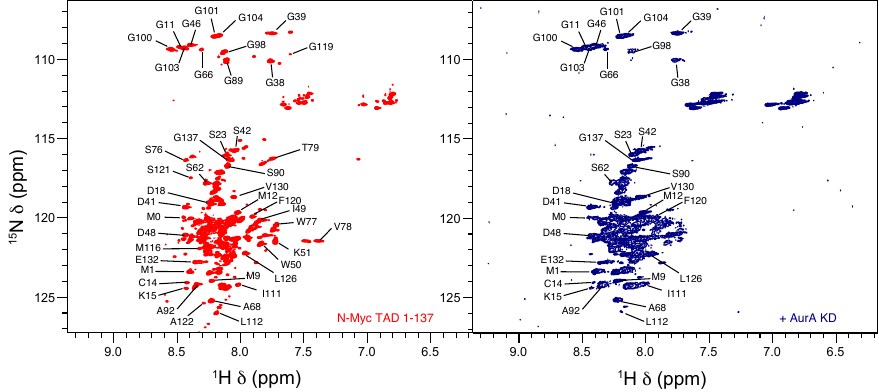

**Figure S5.** ^1^H–^15^N HSQC spectrum for N-myc TAD before (red) and after addition of Aurora A kinase domain (navy blue). Specific peak loss is observed but there was significant attenuation of spectral quality.

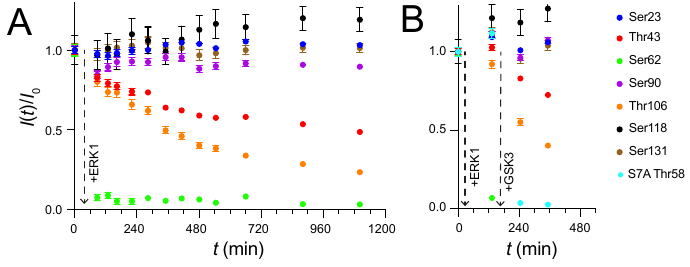

**Figure S6.** (A) Tracking the kinetics of different sites of phosphorylation by ERK1 within N-myc TAD. Rapid and complete phosphorylation of Ser62 is achieved; much slower phosphorylation of Thr43 and Thr106 is also observed. (B) Tracking the kinetics of different sites of phosphorylation by ERK1 and GSK3 within N-myc TAD^S7A^. Rapid and complete phosphorylation of Ser62 by ERK1 is achieved followed by rapid and complete phosphorylation of Thr58; Thr58 does not appear to be significantly affected by ERK1; again slow phosphorylation of Thr43 and Thr106 (by ERK1) is observed.

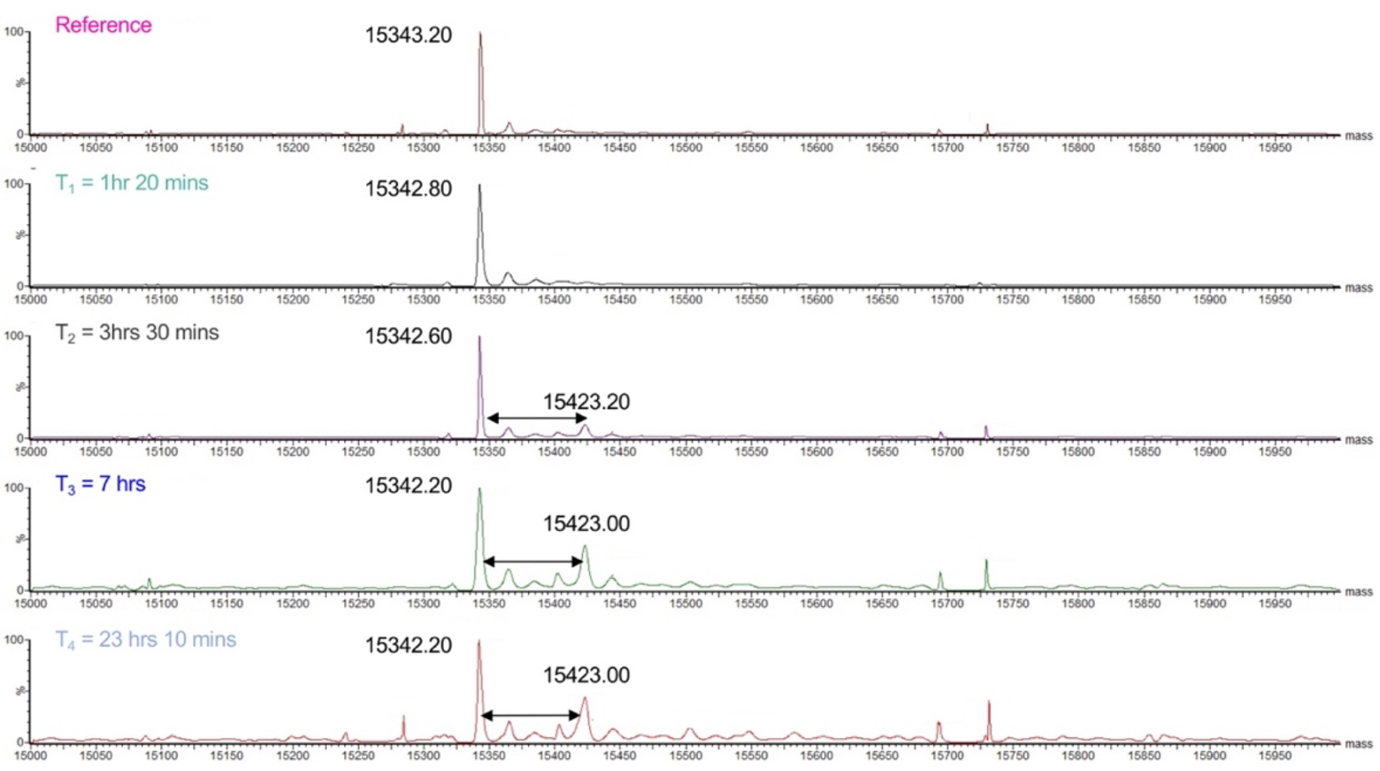

**Figure S7.** Phosphorylation of Thr58 by GSK3 requires priming phosphorylation at Ser62. Intact mass spectra of N-myc TAD were recorded before and at different timepoints after incubation with GSK3.

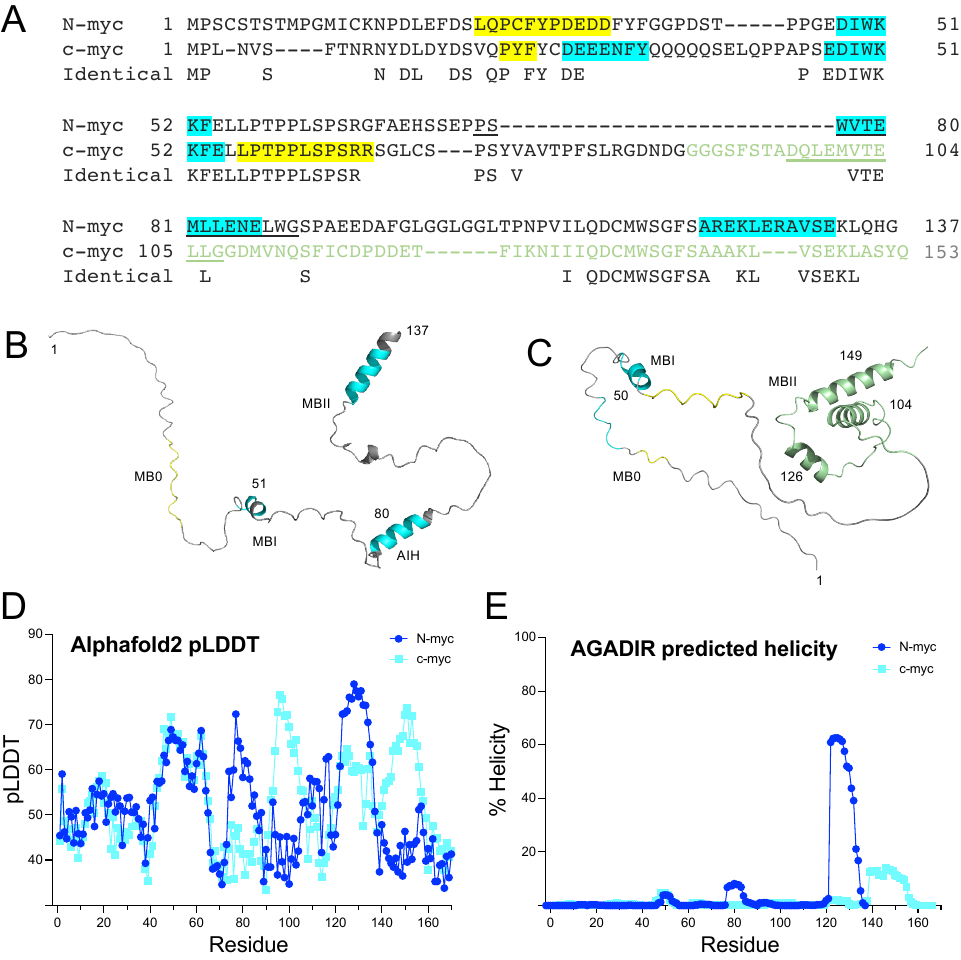

**Figure S8.** Comparison of secondary structure predictions for N-myc and c-myc. (A) Sequence alignment of TADs from N-myc and c-myc, color-coded to show helical regions (cyan) and beta-turn/strand regions (yellow). The region of c-myc TAD for which structural NMR data are unavailable is shown in light green. Underlined sequences are helices in the crystal structures of these regions of N-myc and c-myc. (C) Structure of N-myc based on Alphafold2 prediction, color-coded as (A). (D) Structure of c-myc based on Alphafold2 prediction, color-coded as (A). (D) Residue-specific pLDDT values from Alphafold2 structure predictions for N-myc and c-myc. (E) Output from AGADIR prediction of helical propensity for N-myc and c-myc.
